# Transposable elements hitchhike on Starships across fungal genomes

**DOI:** 10.1101/2025.06.18.660325

**Authors:** Hanne Griem-Krey, Júlia de Fraga Sant’Ana, Ursula Oggenfuss, Yohana Porto Calegari-Alves, Ana Luiza Marques, Markus Berger, Lucélia Santi, Walter O. Beys-da-Silva, Michael Habig

**Author notes:** Corresponding author:* Michael Habig, Walter O. Beys-da-Silva.

## Abstract

Horizontal transfer (HT) of transposable elements (TEs) is a widespread phenomenon in eukaryotes and is often associated with bursts of TE activity. This process profoundly influences genome evolution by introducing novel genetic material and driving genetic variation. However, the precise mechanisms facilitating these transfers remain largely uncharacterized. Here, we report a recent TE burst in the insect-pathogenic fungus *Metarhizium anisopliae*. Our analysis reveals that the actively transposing TEs were introduced via hitchhiking on a so-called Starship—a class of large, themselves horizontally transferable transposons found within the fungal subphylum *Pezizomycotina*. This particular Starship carried 73 TEs, 43 of which exhibited increased copy numbers in the recipient genome, including 508 perfect copies. This expansion triggered extensive structural reshuffling across all chromosomes and led to the formation of a novel chromosome. Remarkably, this structural reorganization was associated with a dramatic phenotypic shift: the loss of pathogenicity. Expanding our analysis to other fungi, we found that Starship-mediated horizontal transfer of TEs is a general phenomenon. A majority (74%) of 618 published Starships also harbor TEs, which can constitute up to 72% of their content. Moreover, Starships serve as sources of actively transposing TEs: 16% of Starships carried at least one TE with a perfect copy found elsewhere in the genome, and identical TEs were observed on Starships from different species. Collectively, our results establish Starships as major vectors of horizontal TE transfer within *Pezizomycotina* and further highlight their profound impact on recipient fungal genomes through TE piggybacking.

## Introduction

Transposable elements (TEs) play a critical role in shaping genomes, contributing significantly to their evolution and the adaptation of species. TEs are classified into two major classes based on their transposition mechanisms: Class I retrotransposons, which transpose via an RNA intermediate, and Class II DNA transposons, which move through a DNA intermediate (1). As mobile genetic elements, TEs can dramatically alter genome architecture, influence gene expression, and create genomic plasticity that enables species to adapt to changing environments. TEs can drive structural genome variation not only via insertions into different sites in different individuals but also through processes such as homologous recombination between TE copies (2), leading to chromosomal rearrangements including translocations, inversions, duplications, and deletions (3–5). For example, *Styx* (also known as REP9), a Class II transposon in the fungal wheat pathogen *Zymoseptoria tritici*, has been independently activated multiple times in distinct populations and is associated with structural variations, triggering several chromosomal rearrangements, including deletions, duplications, and chromosomal fusions (6–8). Transposition events may also result in insertional mutations (9, 10), alter gene expression, or create new protein functions through exon shuffling and TE domestication (11–15). Additionally, TEs serve as sources of regulatory elements (16), spreading cis-regulatory sequences across the genome and reconfiguring gene regulatory networks (17). TE expansions are often associated with structural variant (SV) accumulation, as seen in domesticated crops such as tomato, rice, wheat, and soybean (18–21). Notably, TEs and SVs tend to cluster in certain genomic regions, although it remains unclear whether this colocalization is causal or simply reflects shared tolerance to mutagenic change. To control the potentially harmful effects of TEs, host genomes have evolved various defense mechanisms, including DNA methylation (22), histone modification and heterochromatin formation (23), RNA-silencing pathways (24), and in fungi, repeat-induced point mutation (RIP), which introduces high C-to-T mutation pressures in repetitive sequences to inactivate TEs (25). The vast majority of TEs are considered transpositionally inactive, possibly due to these defense mechanisms, which have been theoretically proposed to ultimately eliminate TEs from most species—a paradox, given their persistent and widespread presence across many genomes (26).

Horizontal transfer of transposable elements (HTT) is increasingly recognized as a significant driver of genome evolution in eukaryotes and may enable TEs to escape the host genome’s silencing mechanisms (26). While most documented cases have focused on plants (27, 28) and animals (29–31), evidence for HTT in fungi remains comparatively limited (32). A recent example includes the proposed horizontal transfer of TEs between rice- and wheat-infecting lineages of the fungal phytopathogen *Pyricularia oryzae* (33). HTTs can involve functionally diverse transposons (34), including those capable of introducing new introns (35). Although thousands of HTT events have been identified across eukaryotes (31, 32, 34), the mechanisms underlying these transfers remain poorly understood. Proposed vectors include viruses and extracellular vesicles (36–39); for instance, LINE1 retrotransposons have been experimentally shown to be packaged in extracellular vesicles and transferred between cultured human cells (38), while other TEs have been detected within large DNA viruses infecting arthropods (36). In fungi, additional mechanisms may also facilitate HTT. Parasexuality—a process allowing genetic recombination independent of meiosis—has been observed in several fungal species and could theoretically introduce TEs into new genomic backgrounds (40, 41). Similarly, horizontal chromosome transfer (HCT), documented in various fungal pathogens, could transfer TE-rich chromosomes between strains (42–46). However, experimental confirmation of TE movement via parasexuality or HCT is still sparse. Interactions with other organisms, such as plants or insects, may also provide opportunities for TE transfer, particularly during parasitic or symbiotic associations that involve close physical contact. In summary, although HTT appears to play an important role in fungal genome evolution, the underlying mechanisms remain poorly characterized and merit further investigation.

Starships are a recently characterized group of massive transposable elements exclusive to fungi within the *Pezizomycotina* subphylum (47–50). Ranging from 15 to nearly 700 kilobase pairs in length (51), these elements are mobilized by a conserved tyrosine recombinase gene known as Captain, which contains a DUF3435 domain located at the 5′ end of the element (52). Unlike traditional transposons, Starships frequently carry diverse cargo genes that encode traits conferring ecological and evolutionary advantages, such as virulence factors (e.g., the necrotrophic effector ToxA) (49, 53), resistance to formaldehyde (54) and heavy metals (52), and biosynthetic gene clusters (53, 55). Typically found as single copies in fungal genomes, Starships have been implicated in promoting genome plasticity by contributing to large-scale structural variations, including chromosomal rearrangements and insertions into lineage-specific regions, as observed in species such as *Verticillium dahliae* (56), *Macrophomina phaseolina* (57), and *Aspergillus fumigatus* (58, 59). A defining feature of Starships is their capacity for horizontal gene transfer (HGT) between fungal species, facilitating the rapid acquisition of adaptive traits. This phenomenon has been documented in both plant and human fungal pathogens, where near-identical Starships have been found across distinct taxa and are frequently associated with enhanced virulence or environmental resilience (52, 56, 60–62). Recently, the horizontal transfer of two Starships was experimentally demonstrated through co-culturing experiments, in which strains containing Starships successfully transferred them to conspecific and heterospecific strains lacking the elements (63). Notably, horizontal transfer occurred even between *Paecilomyces variotii* and *Aspergillus fumigatus*, two species that diverged approximately 100 million years ago (63, 64). These findings establish Starships as an active vector for eukaryotic horizontal gene transfer (63).

In summary, although Starships are known to mediate genome rearrangement and horizontal gene transfer, their role in transferring other TEs and the impact of such events on recipient genomes remain largely unexplored since they have only been recently discovered. Here, we use the insect-pathogenic, asexual fungus *Metarhizium anisopliae* (65) as an informative model to investigate the origin and consequences of horizontally transferred TEs. We identified a specific case, in which a large number of TEs were introduced into one strain via a Starship, subsequently expanded, and ultimately led to genome-wide rearrangements and loss of pathogenicity. We subsequently broadened our analysis to additional fungal taxa, revealing that most of the currently known618 Starships carry TEs, many of which include identical TE copies that have expanded within host genomes - even across species boundaries. These findings suggest that Starships are a major vector for the horizontal transfer and expansion of transposable elements in fungi.

## Results

### *Metarhizium anisopliae* strains NE and E6 differ in accessory chromosomes and transposable element (TE) composition

We characterized genome organization of the *M. anisopliae* strains NE and E6 using PacBio HiFi sequencing (see Table S1 for an overview). Sequence analysis yielded assemblies for both strains with chromosome-level resolution, identifying seven confirmed or eight putative core chromosomes, along with one accessory chromosome in E6 and two in NE (Fig. 1A, Fig. S1). All chromosomes in E6 possess telomeres at both ends, while in NE, three chromosomes are capped at both ends, and an additional three have telomeres on only one end (Fig. 1A). Presence/absence polymorphism analysis across five previously published *M. anisopliae* isolates (Fig. S1A) confirmed that chromosomes chrC, chrD, and chrE are accessory. These chromosomes also showed presence/absence variation across 19 published genomes of various *Metarhizium* species (Fig. S1B), further supporting their accessory nature. The identity and size of the accessory chromosomes were validated by pulsed-field gel electrophoresis (PFGE), followed by sequencing of the excised chromosomal bands, which closely matched the sizes determined from the assemblies. In the NE strain, chrC and chrD measured approximately 1.6 Mb and 1.4 Mb, respectively, and a small core chromosome (chr8, ∼2.3 Mb) was present that lacked a corresponding chromosomal band in strain E6, but later was shown to consist of rearranged sequences of E6 core chromosomes (see below). In contrast, E6 harboured chrE (∼1.3 Mb). The genome of NE is 3.6 Mb larger than that of E6, primarily due to the additional accessory chromosome and an expansion of TEs, which increased from 3.3% in E6 to 9.8% in NE (Fig. 1, Table 1). While this expansion is partly explained by the higher TE content on the accessory chromosomes (Fig. 1A, C), elevated TE content was also observed on the core chromosomes (Table S1). Consistent with previous findings (43), the three identified accessory chromosomes exhibited higher TE density and lower gene content. Additionally, genes located on accessory chromosomes displayed significantly different codon usage patterns, suggesting distinct evolutionary histories compared to those on the core chromosomes (66). In summary, the genomes of the two *M. anisopliae* strains NE and E6 show comprehensive structural differences, even though they do belong to the same species.

**Fig. 1:**
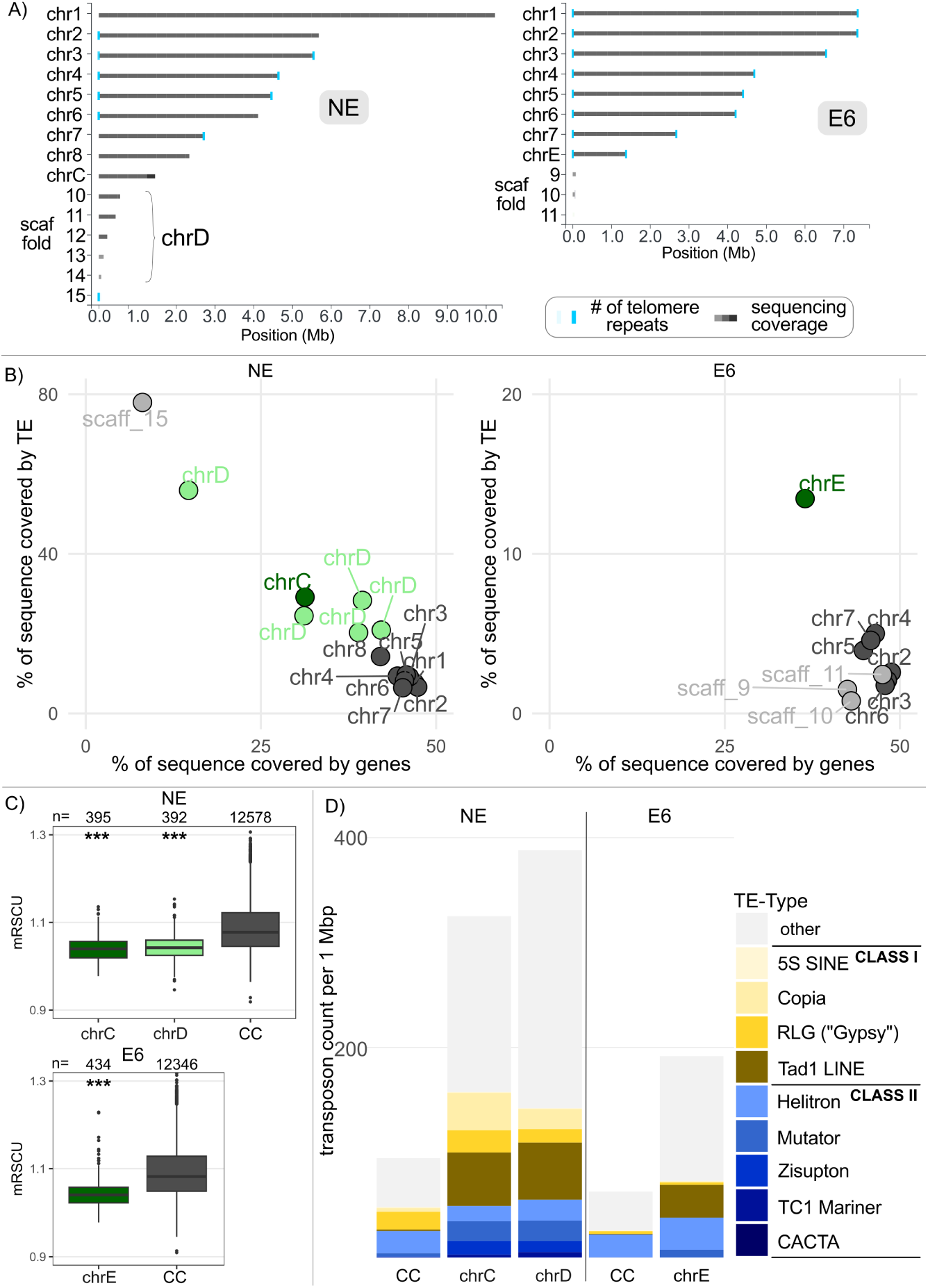
*M. anisopliae* strain NE and E6 differ in genome organization, aaccessory chromosomes and transposable elements. **A)** Graphical representation of the genome assemblies for *M. anisopliae* strain NE (left) and E6 (right). In E6, all chromosomes have been fully assembled from telomere to telomere, and the NE assembly is also at near-chromosome level. **B)** Distribution of genes and transposable elements (TEs) between core and accessory chromosomes for strains NE (left) and E6 (right). A dot plot shows the percentage of the sequence covered by TEs (Y-axis) and genes (X-axis) for core chromosomes (dark grey), accessory chromosomes (light and dark green), and unplaced scaffolds (light grey). Accessory chromosomes are distinct from core chromosomes by having higher TE content and lower gene content. **C)** Genes on accessory chromosomes exhibit a different codon usage bias compared to genes on core chromosomes. The mean gene-wise relative synonymous codon usage (RSCU) is shown for gene transcripts located on accessory and core chromosomes. Statistical significance, as determined by the Wilcoxon rank-sum test, is indicated relative to core chromosomes (CC): *** = p < 0.001. **D)** TEs are overrepresented on accessory chromosomes, and their expansion is more pronounced in NE compared to E6, primarily involving class I (copy-and-paste) transposons

### NE has recently undergone extensive chromosome reshuffling

Synteny analysis of orthologous genes revealed a dramatic loss of macro-synteny between the NE and E6 strains (see Fig. 2A, Fig. 2B). While E6 exhibits a high degree of macro-synteny conservation with chromosome-level assemblies of the more distantly related *M. robertsii* and *M. brunneum*, the comparison between the closely related NE and E6 genomes identified 37 major breakpoint regions (see Table S2A–B). Of these, 28 are present in the fusion of sequences originally located on two different E6 chromosomes (termed “interchromosomal”), while the remaining nine joined non-syntenic regions from the same E6 chromosome, possibly representing large-scale inversions. For example, NE chromosome 1 is a mosaic of sequences syntenic to six different chromosomes in E6, and the newly formed core chromosome chr8 in NE is composed of two large segments syntenic to E6 chromosomes 5 and 7. Based on orthologous gene-based synteny, the breakpoint regions could be localized to intervals ranging from 412 bp to 54,879 bp in length. Phylogenetic analysis of all single-copy orthologous proteins shared between NE and E6 and published *Metarhizium* genomes confirmed that the two strains are closely related and form a monophyletic clade within *M. anisopliae*, along with other published strains (Fig. S2A). Furthermore, no large-scale variation in SNP or small INDEL patterns (a total of 162353 SNPs & small INDELs differentiate E6 from NE) was observed between NE and E6, and the distribution of these variants along the NE chromosomes did not correspond to the mosaic synteny pattern (Fig. S2B). Therefore, we exclude the possibility that the mosaic synteny in NE resulted from a hybridization event between more distantly related species. Instead, we conclude that the genome of NE has recently undergone major chromosomal rearrangements.

**Fig. 2:**
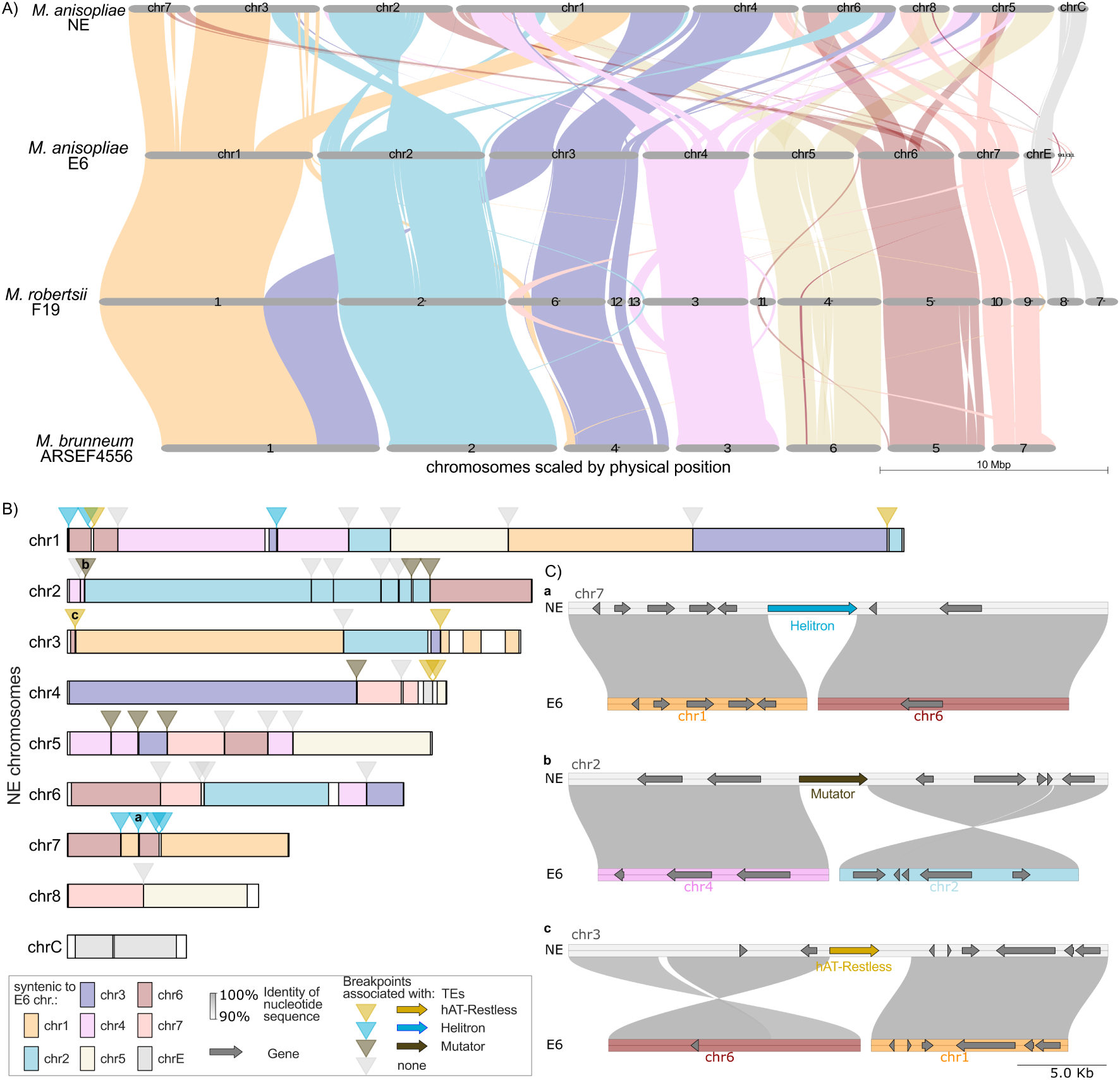
Strain NE has recently undergone extensive structural genome rearrangements, with many breakpoints associated with three distinct transposable elements (TEs). **A)** Macrosynteny between strain E6 and its related species, *M. robertsii* and *M. brunneum*, is highly conserved. In contrast, the NE genome exhibits widespread rearrangements affecting all core chromosomes. **B)** Depiction of chromosomes of NE colored according to the macrosyntenic relationships (based on gene synteny) with E6 chromosomes. Breakpoint regions are marked by triangles, colored to indicate the presence of one of three types of TEs: yellow for hAT-Restless, blue for Helitron, and brown for Mutator. Grey triangles represent breakpoints without these TEs. For clarity, chrD and scaffold_15, which do not contain breakpoints, are not shown. **C)** Representative examples of nucleotide-level synteny at three breakpoint regions (a, b, c as indicated in panel B), each associated with one of the three different TEs. These examples illustrate how TE-associated rearrangements connect sequences that are syntenic to regions on two different E6 chromosomes, which resulted in inter- and intra-chromosomal shuffling. For a complete overview of nucleotide synteny at all breakpoint regions, see Fig. S3. Please note that breakpoint regions associated with Starships are not included.

### Synteny-breaks are frequently associated with three transposable elements (TEs)

In 17 of the identified breakpoint regions, we detected the presence of one of three individual TE sequences that belong to three different TE families: hAT-Restless, Helitron, or Mutator, found in 4, 6, and 7 regions, respectively (see Fig. 2B). In contrast, no TE was detected in the remaining 20 breakpoint regions. Further nucleotide-level alignment of these breakpoint regions allowed us to more precisely pinpoint the synteny breakpoints. In many of the 17 TE-associated breakpoint regions, the TE was on both sides—or at least on one side—directly adjacent to a syntenic alignment (Fig. 2C, Fig. S3A), suggesting that these TEs likely played a role in mediating genome rearrangement at these sites. For the breakpoint regions without an associated TE, we were able to localize the synteny disruption to within 10 base pairs in 17 of the 20 breakpoint regions. Although there was little sequence conservation among these breakpoint-adjacent regions, we observed a potential conserved motif shared among several of them (Fig. S3B–C) – but did not find any sequence similarity to known motifs (TEs or otherwise). Collectively, these results underscore a significant contribution of transposable elements to the formation of synteny breaks, while also indicating that additional, TE-independent mechanisms may underlie genome rearrangement at other sites.

### The TE sequences were horizontally acquired and associated with a recent transposition burst

We detected a total of 196 near-identical copies of the three different TE families associated with the structural rearrangement in the NE genome. The individual copies of each of these three TEs families exhibited very little sequence variation between each other, with the majority being completely identical, indicating a very recent transposition burst (Fig. 3A–B). In most cases, this identity extended to the terminal inverted repeats (TIRs), which are 22 bp and 95 bp in length for the hAT-Restless and Mutator elements, respectively. These TIRs are essential for the transposition of these elements, and their high sequence identity among numerous copies further supports a recent increase in the copy number of these functional transposons.

**Fig. 3:**
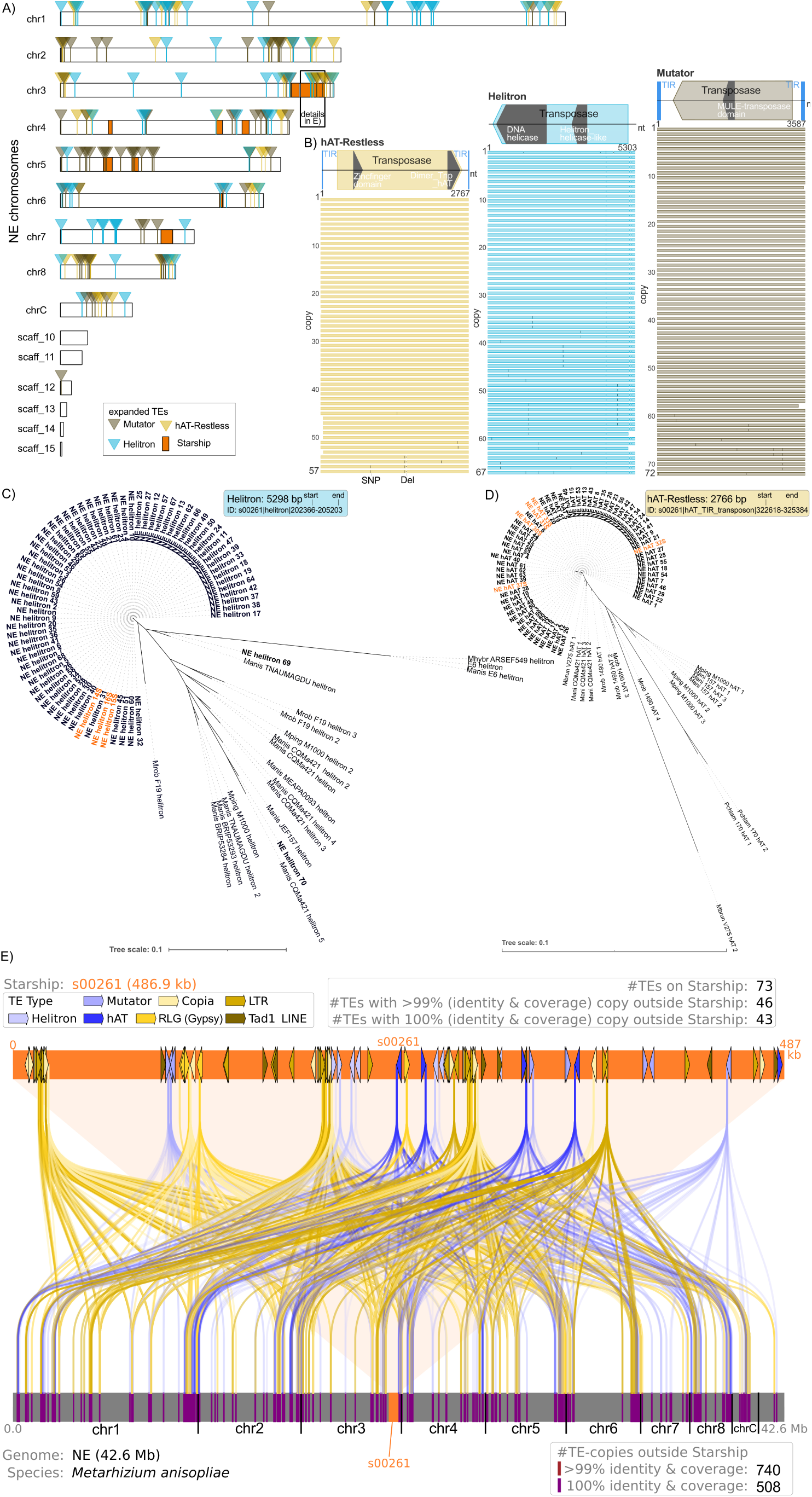
Three transposable elements (TEs) have recently expanded in NE, present in multiple copies, and were likely horizontally acquired via the Starship s00261, along with many additional TEs. **A)** Distribution of near-identical copies of Mutator, Helitron, and hAT-Restless TEs (depicted as yellow, blue, and brown triangles, respectively) across the chromosomes of the NE strain. A total of 196 near-identical copies of these three TEs families are present. Starship regions are highlighted in orange. Copies of all three TE families are located within Starship s00261 on chromosome 3. **B)** Nucleotide alignment of the copies of the three TE families reveals that a large proportion is identical, with only minor differences (e.g., SNPs or small deletions). For orientation, a schematic of their putative structures—including predicted transposase and other functional domains—is shown at the top. **C–D)** Phylogenetic trees of the Helitron (**C**) and hAT-Restless (**D**) TEs based on BLAST hits showing >80% identity and coverage among *Metarhizium* species. Sequences from *M. anisopliae* strain NE are shown in bold, and those located within Starship s00261 are highlighted in orange. Copies of the TE families that recently increased in copy number, including those from s00261, form monophyletic clades distinct from other members of the same TE family or superfamily present in NE. (See also Fig. S4A-D for the phylogeny of the Mutator TE and additional TEs that recently increased in copy number and were located on Starship s00261.) **E)** Detailed map of Starship s00261 (orange, top) with the positions of annotated TEs (including transposon fragments) larger than 100 bp, color-coded by TE superfamily: different shades of yellow for Class I (retrotransposons) and blue for Class II (DNA transposons). At the bottom, location of identical copies (100% identity and coverage) of these TEs found elsewhere in the genome are shown in purple. Connectors between elements are color-coded according to TE superfamily. The numbers above s00261 refer to the individual TEs: it contains 73 TEs >100 bp, of which 46 have at least one copy outside the Starship with >99% identity and coverage. Of these, 43 have at least one fully identical copy, collectively totalling 508 identical copies across the genome. Note: Only 100% identical hits are shown for clarity.

What is the origin of these TEs families that recently increased in copy number? Phylogenetic analysis revealed that these TE families, that increased in copy number, did not originate within strain NE but were acquired – most likely through horizontal transfer. In each case, the TE family members formed a monophyletic group distinct from other members of the same TE superfamily in the NE genome. For example, the copies of the Helitron TE family that increased in copy number and were associated with the breakpoints are clearly distinct from other Helitron TEs present in the NE genome. The phylogenetically closest Helitrons are found in a separate species, *M. robertsii*. Similarly, the copies of the hAT-Restless and the Mutator TEs that increased in copy number and were associated with the breakpoints, were monophyletic and distinct from other hAT-Restless and Mutator elements in the NE genome and most closely related to the respective TE elements in *M. brunneum* or *M. robertsii,* respectively (Fig. 3B, Fig. S4A).

### A horizontally acquired Starship brought along TE sequences that subsequently underwent a burst of transpositions

The presence of three different types of DNA transposons that appear to have been horizontally acquired and contributed to genomic reorganization suggests the horizontal acquisition of larger genetic elements. Consequently, we tested for the presence of active Starships in the NE and E6 strains, as well as in 19 isolates from different *Metarhizium* species. We found that species within the *Metarhizium* genus harbor a high number of Starships (a total of 39) and associated tyrosine recombinase (YR) proteins (Fig. S3 & Table S3). *M. anisopliae* strain NE contains a total of nine Starships, spanning 1.6 Mb in total, and encoding 19 putative tyrosine recombinase (YR) proteins (Fig. S4A–B). These Starships in NE belong to four different families and are absent in *M. anisopliae* E6, which lacks transposed Starships but contains six putative YR genes (Fig. S4B). Phylogenetic and synteny analyses of the Captains (putative YR genes, possibly involved in the excision and mobilization of the Starship) and the associated Starships in NE revealed that the Captains are phylogenetically diverse and distinct from the YR genes present in the E6 strain (Fig S5 D). Interestingly, the Captains of four Starships in NE (s00260, s00265, s00266, s00268) exhibit high sequence similarity, which also extends to portions of their cargo (Fig S4E). However, the insertion sites of these four Starships show no synteny, indicating that the multiple similar Starships found in NE are the result of independent insertion events, rather than the amplification of a single ancestral element during genome reorganization. In conclusion, *Metarhizium* spp. can harbor multiple Starships, with up to nine transposed elements found in a single genome (*M. anisopliae* strain NE). Notably, NE appears to have undergone a recent expansion of Starships possibly through repeated horizontal acquisition events.

One of the horizontally acquired Starships in *M. anisopliae* strain NE brought along multiple transposable elements (TEs) that subsequently multiplied throughout the genome. We observed that Starship (ID=s00261), located on chromosome 3 in NE, contains several copies of the TEs families mentioned earlier, that had recently increased in copy number — namely, three copies of the Helitron family, four copies of the hAT-Restless family, and two copies of the Mutator family. This led us to hypothesize that these three TE families were introduced into NE via horizontal transfer of this Starship. Upon examining all 73 distinct TEs (it is important to note that these encompass full-length TEs as well as fragments greater than 100 bp, which may include solo-LTRs or products of nested insertions, for instance - all of which will subsequently be referred to as TEs) located on Starship s00261, we found that the majority (46) had at least one copy elsewhere in the genome with >99% sequence identity and >99% coverage. Of these, 43 TEs had at least one identical copy (100% sequence identity across 100% of their length) outside the Starship with 21 of these 43 being longer than 1000 bp (Table S4). These TEs include both DNA transposons and retrotransposons. In total, 740 copies (or 508 at 100% identity) of TEs originally located on Starship s00261 were now dispersed throughout the NE genome (Fig. 3E). Assuming a random distribution of the 100% identical TEs across the genome, their density within the Starship s00261 is significantly elevated (p = 1 × 10⁻^20^, Fisher’s exact test with Bonferroni correction). This suggests that either the Starship serves as a preferential integration site for these actively expanding TEs, or it is the source from which they spread. Further phylogenetic analysis of three highly expanded, larger (>1,000 bp) TEs on Starship s00261 — two distinct RLG, formerly known as “Gypsy” (67), and one LTR element — revealed that the expanded copies cluster together phylogenetically and are distinct from other members of the same families or superfamily elsewhere in the genome. Both RLG (Gypsy) retrotransposons have other members of the same TE family in in the NE genome, but their closest phylogenetic relatives are found in *M. pinghaense* or *M. robertsii* (see Fig. S4B, D). In the case of the LTR retrotransposon, no close relative was found within the *Metarhizium* genus, hence we looked for closely related sequences beyond the *Metarhizium* genus and found the most closely related sequence in *Hirsutella rhossiliensis* (see Fig. S4C), a soil-associated ascomycete that infects nematodes (68). Hence all large TEs that increased recently in copy number in NE and were locate on Starship s00261, are phylogenetically separate and hence most likely have been horizontally acquired. These are also significantly enriched on the Starship, therefore support the conclusion that Starship s00261 introduced these TEs into NE, from where they subsequently increased in copy number and invaded the genome which in turn resulted in the complete reorganization of the genome.

### Structural reorganization of the NE genome is associated with reduced pathogenicity

To assess the phenotypic impact, we compared the pathogenicity of strain NE to strain E6 in Asian blue tick, *Rhipicephalus microplus* an economically important tick of mammal livestock that can be controlled through applications of *M. anisopliae* as a biocontrol agent (Fig. 4A). Strain NE exhibited significantly lower pathogenicity than E6, being not different from the control treatment, indicating that, unlike E6, NE is unable to kill *R. microplus*. The reduced pathogenicity in NE compared to E6 was not restricted to *R. microplus*. In a previous study *M. anisopliae* strain NE was less infective than the E6 strain against the cotton stainer bug *Dysdercus peruvianus*, an insect pest causing heavy losses in cotton plantations (69). NE had also a reduced sporulation capacity compared to E6 (69). Taken together these earlier and our results here indicate that NE has a decreased pathogenicity. Therefore, we tested whether the reduced pathogenicity of NE could have been a consequence of the genome reshuffling. During early infection, enzymatic activity in the fungal secretome plays a critical role in cuticle degradation (70–73) and we identified the 48h differential mycelial secretome, induced *in vitro* by liquid culture supplemented with tick cuticle. In the secretomes we determined the activity of six infection-related enzymes. Five of the six tested enzymes showed reduced activity in NE, while PR2 activity was increased, suggesting that strain-specific differences potentially emerge early in infection and may contribute to the observed variation in pathogenicity (Fig. 4B). Proteomic analysis of the secretome further revealed major differences in its composition: 228 proteins were more abundant in E6, 16 in NE, and 51 showed similar levels between the strains (Fig. 4C, Table S5), highlighting a pronounced divergence in secretome profiles induced by host components. To test whether the structural variation of the genome might have caused these differences we then assessed whether genes encoding the secreted proteins were located near structural variations (expanded TEs, breakpoints, or Starship elements). Although a greater proportion of differentially expressed genes (53 of 244) were located within 10 kb of such variations compared to similarly expressed ones (5 of 51), this trend was not statistically significant (Fisher’s exact test, *p* = 0.07763). For some important individual genes however, there appeared be a correlation. Of particular interest is a subtilisin-like protease, PR1C—a key pathogenicity factor potentially contributing to host specificity (72, 74, 75). In strain E6, the gene encoding PR1C is located on accessory chromosome chrE (geneID: g11457), whereas in NE, it has been relocated to chr4, near a copy of the hAT-Restless TE - that had recently increased in copy number - and a breakpoint where regions syntenic to chrE and chr5 in strain E6 have been fused (Fig. 4D, geneID: g7795). PR1C showed reduced expression in the secretome of NE compared to E6, which is also reflected by the lower detected enzyme activity for PR1 (Fig. 4B). Furthermore, a neighbouring gene (E6 geneID: g11461) encoding a putative galactose oxidase precursor, located near the PR1C-encoding gene in E6 was also relocated by the structural variation in NE (NE geneID: g7793) and similarly showed reduced protein expression in the NE compared to the E6 secretome (Fig. 4C). In addition, a putative biosynthetic cluster for the insecticidal metabolite destruxin (76) was modified and several additional biosynthetic genes have been lost from the cluster and were also not present in the rest of the genome, possibly due to an insertion of a copy of the Helitron TE family in the NE strain (Fig. 4D). Taken together, these findings suggest that structural variation in NE may have altered the local genomic environment, leading to altered expression of key secreted proteins or insecticidal compounds and potentially explaining the strain’s loss of virulence and pathogenicity.

**Fig. 4:**
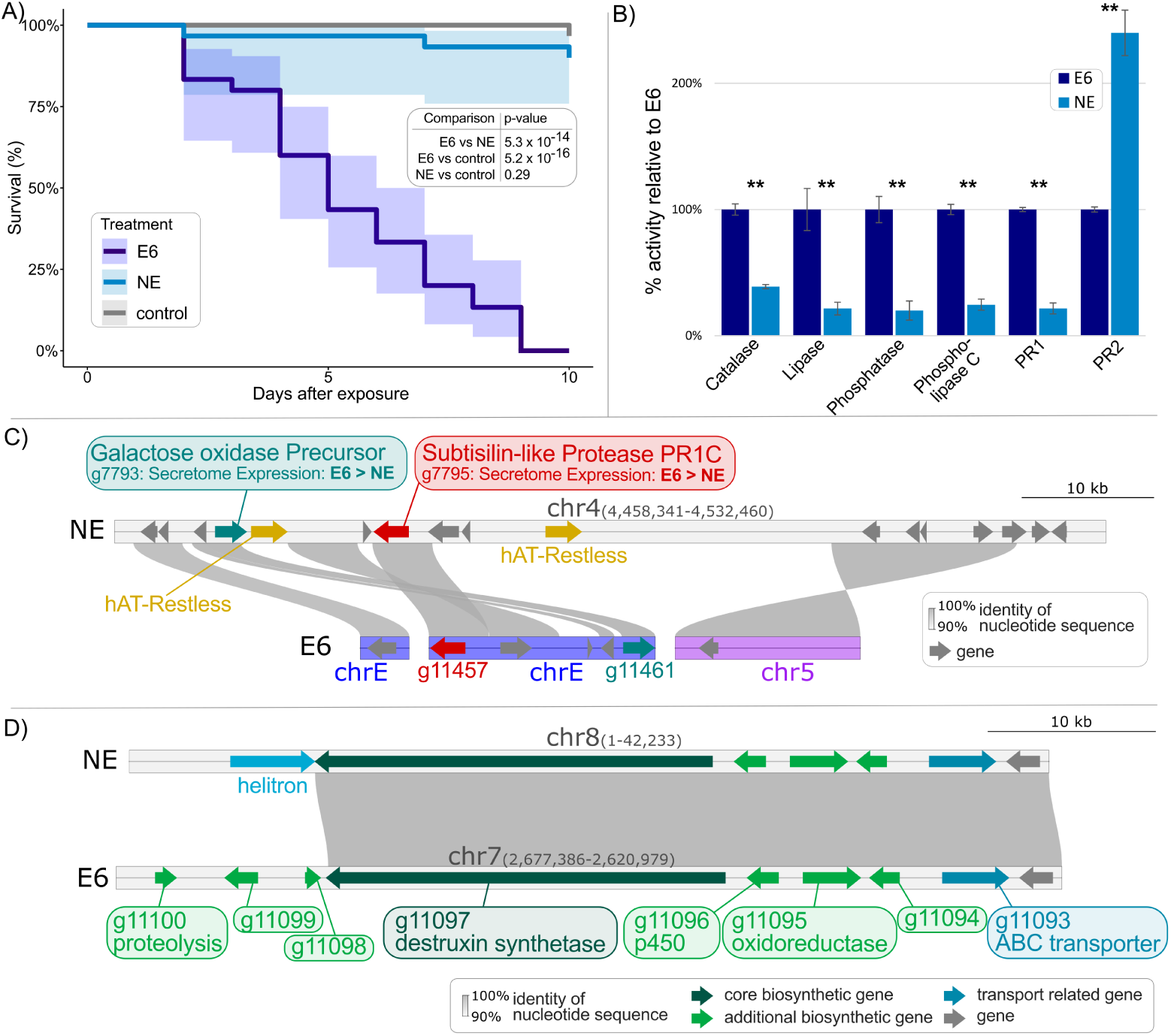
Effect of structural variation on the pathogenicity and secretome of *M. anisopliae* strain NE. **A)** Kaplan-Meier survival curves of *Rhipicephalus microplus* ticks inoculated with *M. anisopliae* strain NE (light blue), strain E6 (dark blue), or a control (grey). Strain NE is less pathogenic than strain E6 and does not differ significantly from the control group. Shaded areas are 95% confidence intervals of the mean survival. For each treatment, the data of three independent replicates with ten individuals each were pooled. Statistical significance was determined using the log-rank test with BH-adjustment for multiple testing. **B)** Comparison of enzymatic activity between strains E6 and NE. Activities of each enzyme class were normalized to those of strain E6. Catalase, lipase, phosphatase, phospholipase C, and PR1 exhibited significantly reduced activity in strain NE, while PR2 activity was increased compared to E6. Statistical significance of differences between E6 and NE, as determined by an unpaired t-test, is indicated above each bar (** = p < 0.001). Data are expressed as the mean of three replicates ± standard error. **C)** Genomic context of two genes—g7793 (encoding a putative galactose oxidase precursor) and g7795 (encoding a putative subtilisin-like protease PR1C)—is shown in the figure. Their surrounding regions have been extensively remodeled in NE (upper panel) compared to their syntenic regions in E6 (lower panel), where orthologous genes are shown in the same color. The presence of two copies of the hAT-Restless TE and a nearby breakpoint (within 10 kb) in NE may have contributed to reduced expression of these genes. Note: Only TEs that had recently increased in copy number are shown for clarity. D) Possible effect of an integration of a TE of the Helitron family on the organization of the putative destruxin A cluster. The destruxin synthase gene as the core biosynthetic gene (dark green) and several additional biosynthetic genes (light green) are present in both E6 and NE. The additional biosynthetic genes g11098, g11099 and g11100 have been lost from the cluster (and from the genome) in NE, possibly due to the insertion of a copy of this Helitron TE.

### TEs were frequently found on Starships and often had additional copies in the rest of the genome

We next broadened our analysis to other fungal taxa and investigated whether TEs are generally prevalent on Starships and, if so, whether these Starship-associated TEs also have copies in the remainder of the genome. To this end, we analysed 618 published and unique Starships available in the Starbase repository (77). We included the 39 Starships identified in the Metarhizium genus in this study, thereby including starships from a total of 164 distinct species, and annotated all known TE families using EDTA (78). The majority of Starships contained known TEs, which in some cases accounted for more than 50% of the sequence, with a maximum of 71.8% TE coverage observed in Starship SBS000643 from *Aspergillus chevalieri* (Fig. 5A, Table S4). In addition to *Aspergillus*, high TE contents (>30%) were also observed in Starships from the fungal genera *Histoplasma*, *Alternaria*, *Metarhizium*, *Lassallia*, and *Pyricularia* (Table S4). Where accessions of the genomes with the Starships were available, we assessed whether TEs identified on Starships had either highly similar copies (>99% sequence identity and >99% coverage), or perfect copies (100% sequence identity and 100% coverage) elsewhere in the genome. Of the 422 Starships for which this analysis was possible, 314 contained at least one TE. Of the 314 Starships, 31% (97) and 21% (67) had a highly similar or a perfect copy of a TE elsewhere in the genome, respectively. Of the latter, 35 consisted of TEs longer than 1000 bp (Table S4). A higher proportion of Starships containing TEs with perfect copies was observed in species of *Aspergillus*, *Metarhizium*, *Alternaria*, *Colletotrichum*, and particularly *Pyricularia*, where all five of the six described Starships from two different genomes contained at least one TE with a perfect copy elsewhere in the genome (Fig. 5B, Table S4). Hence, we conclude that there has been a recent exchange of TEs between Starships and their surrounding genomes. Either the TEs originated in the genome and transposed into the Starship, or vice versa.

**Fig. 5:**
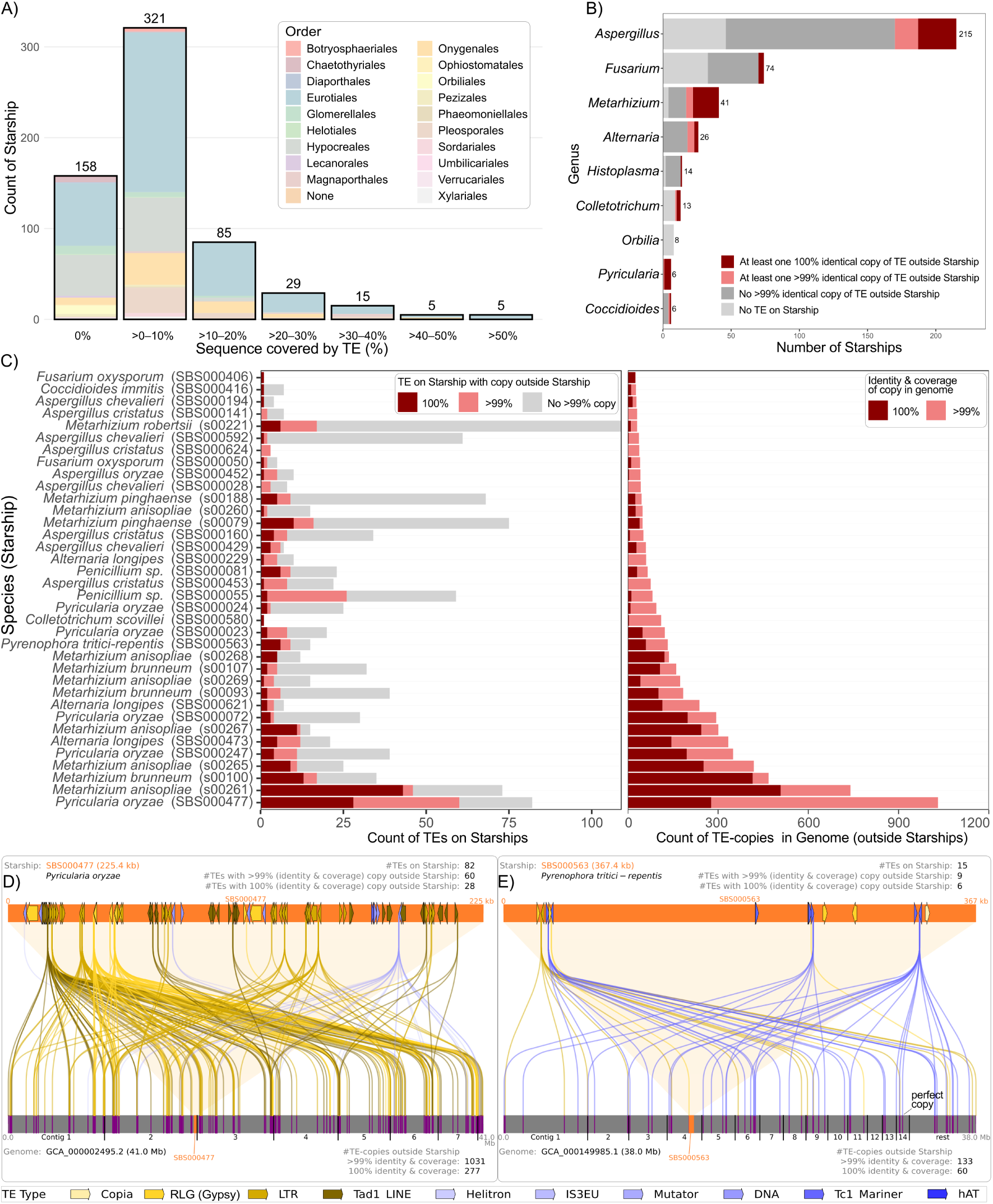
The majority of Starships contain transposable elements (TEs), many of which have multiple identical copies elsewhere in the genome. **A)** Binned percentages of Starships covered by known TE families, colored according to the taxonomic order of the species in which they were first described. Most Starships contain known TE families, with 59 Starships having more than 20% TE content. **B)** Counts of Starships—limited to those with genomic data available—categorized by TE content: Starships without TEs (light grey), Starships containing TEs but no genomic copies outside the Starship with more than 99% sequence identity and coverage (light red), and Starships containing TEs with identical genomic copies elsewhere (red). *Aspergillus*, *Metarhizium*, and *Pyricularia*, in particular, show a high fraction of Starships harboring at least one identical TE copy outside the Starship. Only genera with at least five Starships and available genomic data are shown. **C)** Detailed analysis of individual Starships with at least 25 TE copies outside the Starship. Left panel: Number of TEs on Starships, distinguishing between those TEs with at least one external genomic copy at >99% sequence identity and coverage (light red) and those with at least one identical genomic copy (dark red). Many Starships harbor multiple TEs with highly similar or identical genomic copies. Right panel: Number of TE copies for these Starships, with several Starships showing high numbers of very similar or identical copies elsewhere in the genome. Panels **(D)** and **(E)** show graphical representations of TEs on two example Starships (orange bars at top) and their perfect genomic copies outside the Starship (purple markers on continuous grey genome bars below): SBS000477 in *Pyricularia oryzae* **(D)** and SBS000563 in *Pyrenophora tritici-repentis* **(E)**. TE families are color-coded: Retrotransposons appear in different shades of yellow, while DNA transposons are shown in various shades of blue. Please note that, for clarity, contigs/chromosomes or scaffolds of the genome assembly were ordered and renamed according to decreasing size. Only contigs longer than 1 Mb are shown individually, while all contigs shorter than 1 Mb are grouped together at the right end.

Many Starships contained more than one TE with a perfect copy elsewhere in the genome in which they reside in, which would be more indicative of the Starship being the source of the TEs. The maximum number of different TEs on a single Starship with perfect copies elsewhere in the genome was 46 and observed in the previously described s00261 in *M. anisopliae* strain NE, with 28 in Starship SBS000477 of *Pyricularia oryzae* (see Fig. 5D), 13 in s00100 of *M. brunneum*, and six (out of a total of 15 TEs) in SBS000563 of *Pyrenophora tritici-repentis* (Fig. 5C, Table S4). Again, the number of very similar (>99% sequence identity with >99% coverage) or identical (100% sequence identity and coverage) TE copies elsewhere in the genome can be very high. For example, in Starship SBS000477 from *P. oryzae*, 28 TEs that have at least one perfect copy outside the Starship have a total of 152 perfect copies elsewhere in the genome, along with 989 very similar copies. A Fisher’s exact test assessing whether identical TE copies are randomly distributed across the genome found that 24 of the 67 Starships were significantly enriched for these. Combined with the fact that, in many cases, multiple TEs from different TE classes were involved, we conclude that Starships are the most likely source of these recently actively transposed TEs.

### Starships from different species contain identical TE copies indicating HTT between species

We next asked whether different Starships—either from the same or different species—can share identical TEs, which would further support the idea that Starships can shuttle TEs between species. To test this, we searched for perfect copies of TEs (100% sequence identity and 100% coverage) located on different Starships. Among the 467 Starships with available species-level information, nearly 50% (223) harbored at least one TE that had a perfect copy on another Starship in the same or in a different species. This phenomenon was particularly common in Starships from the genera *Fusarium*, *Penicillium*, *Alternaria*, *Aspergillus*, *Metarhizium*, and *Pyricularia* (Fig. 6A, Table S6). Remarkably, 18.6% (87) of all Starships contained at least one TE with a perfect copy on a Starship from a different species, of which in 17 Starships at least one shared identical TE was longer than 1000bp (Table S6). Tad1 LINE elements appear especially prone to being shared between different Starships, both within and across species. One striking example is a Tad1 LINE element of 1,823 bp that is nearly identical in several Starships of *A. oryzae* (e.g., SBS000219), as well as in Starship SBS504 of *A. flavus*. Similar copies are also shared between *A. oryzae* and *A. sojae* (e.g., SBS00258 and SBS00342) (Fig. 6B). We found that identical TEs are almost exclusively shared between Starships from species within the same genus, and that some genera are more prone to this sharing than others. In the genus *Aspergillus*—which currently has the highest number of annotated Starships—identical TEs are shared between Starships from 13 different species, with the highest number of shared elements found between *A. flavus* and *A. oryzae* (Fig. S7A). In *Metarhizium*, identical TEs are shared among five species, with the most frequent sharing observed between *M. anisopliae* and *M. brunneum* (Fig. S7B). In the genus *Penicillium*, identical TEs are shared between three species, most prominently between *P. chrysogenum* and *P. griseoroseum* (Fig. S7C).

**Fig. 6:**
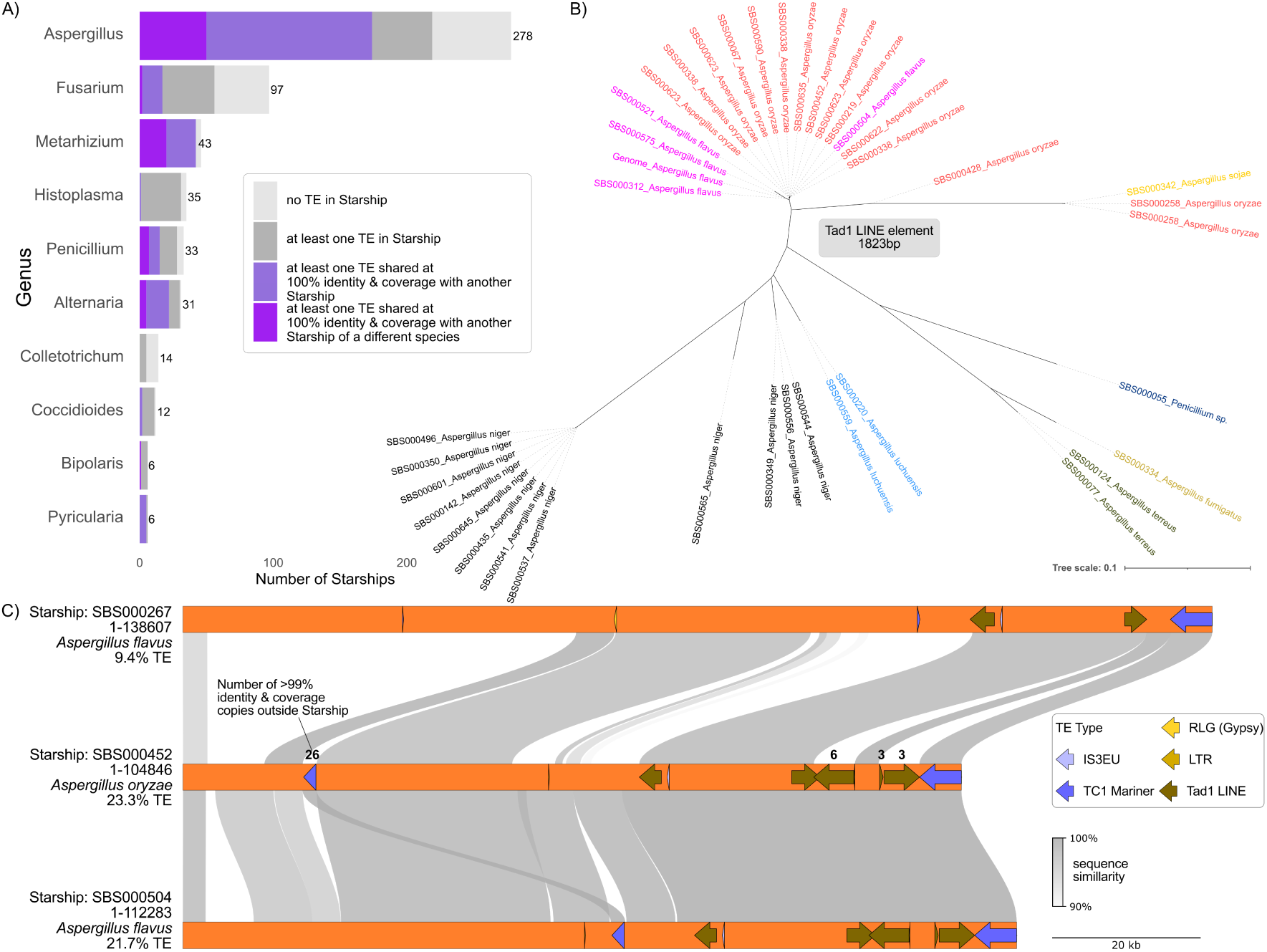
Transposable elements (TEs) located on Starships may have identical copies on Starships from different species and appear to have expanded in the genome. **A)** The number of Starships per genus containing at least one TE with a perfect copy (100% sequence identity and 100% coverage) in another Starship from either a different species (dark purple) or the same species (light purple) is shown. Starships containing TEs without such identical copies are indicated in dark grey, while those lacking TEs entirely are shown in light grey. Genera such as *Aspergillus*, *Metarhizium*, and *Penicillium* exhibit a high proportion of Starships harboring TEs shared between species, suggesting horizontal transfer. Only Starships with associated phylogenetic information at the species level were included in the analysis. **B)** A phylogenetic tree showing an example Tad1 LINE retrotransposon (1832 bp) with identical copies present on Starships from both the same and different species, with species distinguished by color. **C)** Synteny analysis of three different Starships (in orange) from two species (*A. flavus* and *A. sojae*); arrows indicate TEs, while genes have been omitted for clarity. TEs shared between species via Starships have expanded in *A. oryzae*. Connections are shaded in different shades of grey based on nucleotide sequence identity. TE families are color-coded: Retrotransposons appear in different shades of yellow, while DNA transposons are shown in various shades of blue.

Starships could act as both the source and destination of transposing transposable elements (TEs), facilitating their movement between species. We find evidence for both i) TEs located on Starships that have been horizontally transferred between species and subsequently increased in copy number in the recipient genome, and ii) TEs that are actively increasing in copy number within the genome integrating into Starships, potentially setting the stage for future horizontal transmission. Between *A. oryzae* and *A. flavus*, three Starships share syntenic regions that include TEs, indicating horizontal transfer (Fig. 6C). In *A. oryzae* specifically, four of these TEs—one Tc1/Mariner and three Tad1 LINE retrotransposons—have recently expanded and show very high sequence identity and coverage (>99%), making post-transfer increase in copy number the most parsimonious explanation. Similarly, we observe syntenic Starships shared between *M. brunneum* and *M. pinghaense* that include TEs (Fig. S8A), again indicating horizontal transfer. Although the direction of transfer is unknown, in *M. brunneum*, a Tad1 LINE retrotransposon (along with other retrotransposons) appears to have actively transposed and increased in copy number and likely invaded additional Starships. In this case, Starships act not only as vehicles for horizontal transfer but also as both source and destination for actively transposing TEs. Another example is shown in Fig. S8B, where four Starships from two distinct *Aspergillus* species (*A. oryzae* and *A. sojae*) share synteny that includes six TEs, indicating horizontal transfer of those elements. On two of the Starships, numerous copies of TEs—known to have increased in copy number elsewhere in the genome—are integrated in positions that break synteny, suggesting recent insertion events after the horizontal transfer of the Starship. Here, the Starships most likely served as targets for actively transposing TEs. In conclusion, we find evidence of reciprocal exchange between Starships and the rest of the genome, enabling the horizontal transfer of TEs within and across fungal taxa.

## Discussion

Here, we show that in *M. anisopliae*, a large number of TEs were introduced into the NE strain via a Starship. These TEs subsequently transposed actively, dramatically increasing their copy number, and ultimately contributed to genome-wide rearrangements and loss of pathogenicity. Our broad-scale comparative analysis of 618 available Starships across the *Pezizomycotina* subphylum further revealed that the majority carry TEs. Many of these TEs have identical copies within the rest of the genomes, and importantly, identical or very similar copies of these TEs are also found on other Starships, even across species boundaries. This presence of identical or very similar copies on Starships across species strongly suggests that Starship are important facilitators of horizontal transfer of TEs. The high similarity between these individual TE copies further indicates that these horizontal transfers must have been very recent.

Are Starships the source or rather a target of TEs? When accounting for the relative size of Starships within the genome, we observed that 24 of the 67 Starships containing at least one TE with a perfect genomic copy exhibited a significantly higher TE density inside the Starship than in the surrounding genome. This suggests that Starships are more likely to act as the source rather than the recipient of these TEs. Moreover, we found identical TEs on Starships isolated from different species, for some of which the TEs recently must have increased in copy number. This supports the conclusion that Starships are not only a source of expanded TEs but also act as vectors that shuttle them between species within the subphylum Pezizomycotina. Therefore, Starships should be recognized not only as vehicles of frequent horizontal gene transfer (HGT) (63) but also as facilitators of horizontal transposon transfer (HTT), complementing previously reported HTT mediators such as extracellular vesicles and viruses (36–38). Certain genera—*Aspergillus*, *Metarhizium*, *Penicillium*, *Pyricularia*, and *Fusarium*—appear particularly susceptible to Starship-mediated HTT, although this observation may be biased by non-random sampling in Starship detection studies. TEs are predominantly shared among species within the same genus, although in some cases, identical TEs were found on Starships across different genera. This mirrors the observed pattern of more frequent Starship transfer among identical or closely related species compared to distantly related ones (63). We note that many of the TEs shared between starships and their host genomes, as well as between starships in different species, are likely fragments due to their short length (>100 bp). However, we observed similar results for longer TEs (>1000 bp), more likely representing full length TEs, which supports our conclusion that Starships are facilitators of HTT. We also note that by using a very conservative threshold of >99% identity for individual TE elements, we have likely missed older Starship-mediated HTT events, as well as newer ones that are affected by an increased mutation rate due to RIP. Consequently, the contribution of Starships to the spread of TEs between species may be even more widespread than our analysis indicates.

An alternative explanation for the enrichment of Starships with expanded TEs is that TEs may preferentially transpose into Starships—although the underlying mechanism remains unclear. Some Starships are known to integrate into AT-rich regions, 5S rDNA loci, or even directly into other TEs (79), which are also known as preferential insertion sites for TEs. This may result in Starships and active TEs occurring in close proximity to each other. In addition, TE insertions also co-localize with open chromatin (80), hence if the chromatin state of Starships were open, then they might indeed be some of the only regions where active TEs are able to insert. But still, the apparent tendency of TEs to target (active) Starships remains puzzling. Starships are frequently horizontally transferred; for example, in the genus *Paecilomyces*, between 18–27% of Starships show evidence of horizontal transfer under natural conditions (63). This high frequency of horizontal transfer renders Starships conceptually similar to conjugative plasmids in bacteria for horizontal transfer of sequences. It might therefore be possible that active Starships could be preferentially targeted by TEs to increase their chances of horizontal transmission. A precedent for such a mechanism exists in bacteria: the transposon Tn7 is known to specifically insert into conjugative plasmids, likely recognizing them via features associated with lagging-strand DNA replication (81). This strategy enhances the probability of horizontal transfer while minimizing the potential costs of transposition (81). Given that Starships may be associated with circular DNA intermediates, which would be unique and recognizable (50), they could similarly provide an opportunity for TE mobility and spread. While this would represent an intriguing parallel to conjugative plasmids, the high diversity of expanded TEs (from both major TE classes) observed within Starships argues against a single, conserved mechanism. Irrespective of the mechanism, it has been proposed that most actively transposing TEs that are then recognized as TE bursts might enter the genome via Horizontal Transposon Transfer (HTT) (26, 28, 82). This is because defence mechanisms, such as RIP, would require time to recognize and deactivate newly introduced TEs. Hence, the horizontal transfer of TEs might be an integral part of TE life cycles, and the possibility that TEs might exploit Starships for horizontal transfer remains an exciting avenue for future research.

We find that expanded TEs have led to massive structural reorganization of the genome in *M. anisopliae* strain NE. It is striking that only DNA transposons are involved in the structural variation, despite the expansion of several larger (>1,000 bp) retrotransposons in the same strain. Two of these DNA transposons, Mutator and hAT-Restless elements transpose by generating double-strand breaks (DSBs) and form hairpin intermediates either in the flanking regions or within the transposon itself, through a mechanism resembling V(D)J recombination by binding to terminal inverted repeats (83, 84). These DSBs can be repaired replicatively via homologous recombination (HR) using a homologous chromosome, sister chromatid, or another genomic copy as a template— potentially leading to sequence homogenization (85, 86)—or through non-replicative repair via non-homologous end joining (NHEJ) (87). The third TE family associated with breakpoints in NE is a Helitron which replicate via a unique rolling-circle mechanism (peel and paste mechanism) and are capable of capturing and mobilizing host genes (88, 89). Widespread across eukaryotes, Helitrons have significantly influenced genome evolution, with horizontal transfer events contributing to genomic changes in *Xenopus laevis* and bats (90, 91). Although Helitrons can capture adjacent sequences, these smaller elements rarely exceed 10 kb and typically carry only gene fragments rather than full-length coding sequences (88, 92, 93) and therefore this alone cannot account for the widespread chromosomal reorganization we observed. In conclusion, the structural variation observed in *M. anisopliae* strain NE appears to result from a combination of homologous recombination between identical TE sequences and the direct effects of transposition. Mutator and hAT-Restless elements induce DSBs, while Helitrons may introduce single-strand breaks, both increasing the likelihood of recombination. The abundance of identical, likely active TEs may fuel a self-reinforcing, runaway process. However, about half of the breakpoints lack identifiable TEs, and the low sequence conservation in these regions suggests the involvement of additional, as yet unknown, mechanisms. Overall, the extensive structural changes observed in *M. anisopliae* strain NE are at least partially driven by the expansion and activity of Starship-derived DNA transposons.

The observed chromosomal rearrangements could have a major impact on fungal speciation, in this particular case in the taxon *Metarhizium anisopliae*. The burst of TE activity and the resulting extensive chromosomal reorganization in the strain NE, compared to the more conserved E6 strain, may lead to rapid genetic differentiation and reproductive isolation, as previously hypothesized (94). Although meiotic recombination has not yet been documented in *M. anisopliae*, it seems highly unlikely that recombination would be successful between the extensively reshuffled genome of strain NE (or its descendants) and the more ancestral genome of E6. In diploid heterozygotes, such divergence could disrupt chromosomal pairing, crossing over, or segregation during meiosis. As a result, the offspring would likely be inviable (or completely sterile) or suffer from drastically reduced fertility or fitness (reviewed in (95)) and hence *M. anisopliae* strain NE is most likely now reproductively isolated – and could be considered a new species. Additionally, SVs may alter the epigenome, gene regulatory networks, or chromatin organization within the nucleus (96). Such changes could explain the observed differences in the secretome between strains NE and E6, which cannot be entirely attributed to the vicinity of the corresponding genes to TE insertions or breakpoint locations alone. Regardless of its impact on potential speciation, the extreme structural genome reorganization observed in the NE strain is correlated with changes in pathogenicity and life history strategy. Our results indicate that most proteins previously linked to pathogenicity and associated with TEs, breakpoints or Starships (proteases, chitinases, oxidoreductases and other effector proteins) were down-regulated in the NE strain. This provides another compelling example of structural variation as a major driver of genomic and phenotypic diversity, adding to the known cases reported among pathogenic fungi. (97).

In conclusion, we here show that Starships are frequent vehicles for the horizontal transfer of transposable elements, which can subsequently expand and exert substantial effects on recipient organisms. As such, Starships should be recognized as important vectors for the horizontal spread of active transposable elements.

## Methods and Materials

### Strains & Culture

*M. anisopliae* strain E6 and NE were isolated from *Deois flavopicta* and *Mahanarva posticata* insects, respectively. Both strains were isolated from Brazil: E6 from Espírito Santo state and NE from Pernambuco state.

The strains E6 and NE were cultivated for conidia and mycelia production as previously described (98). Briefly, spores (10^6^ mL^−1^) were added to propylene bags containing 100 g of rice and 30 mL of 0.5% peptone, and left at 28 °C for 14 days to promote conidia production. The dry spores were collected for further suspension production. Then, spores (10^7^ mL^−1^) were inoculated in 70 mL of basal medium (0.6% NaNO_3_, 0.2% glucose, 0.2% peptone, and 0.05% yeast extract w/v) encompassing cuticles of *R. microplus* (0.7%) and cholesteryl stearate (0.05%), for triggering the infection machinery (75). The flasks were incubated at 28 °C with shaking (150 rpm) for 48h. After growth, 0.25% (v/v) Triton X-100 was added and mixed to extract enzymes and proteins from the mycelia external surface. Afterwards, the mycelia were harvested by filtration through a Whatman n^o.^ one filter paper. These filtrates containing secreted proteins were frozen and later used for enzymatic assays.

### DNA extraction

For DNA extraction cells were propagated in liquid culture using SBD medium with a final glucose concentration of 8% at 28°C at 200 rpm. Cells were harvested by centrifugation (3200 x g, 20 min at RT and washed twice with H_2_O before, the supernatant removed and the DNA was purified using a modified CTAB protocol with two Phenol:chloroform extractions (99).

### Sequencing and Pulsed-field Gel Electrophoresis

DNA was sequenced at BGI Genomics, Hongkong using PacBio Revio system. Pulsed-field Gel Electrophoresis was conducted and gel bands were extracted and sequenced as described here (43).

### Enzymatic Activities

The enzymatic assays were carried out as previously described (100). All assays were performed in three replicates using culture supernatant prepared as described above. All enzymatic activities were expressed as specific activity (U/mg of protein) based on total protein content measured using the bicinchoninic acid (BCA) protein assay (Pierce, Rockford, USA). The statistical analysis was determined by Unpaired t-test using GraphPad software.

For lipase activity, solution 1 (ρ-nitrophenyl palmitate (ρNPP) (Sigma, USA) 3 mg/mL in isopropanol) was added in droplets to solution 2 (Tris-HCl 50 mM pH 8.0, 0.11% gum arabic, 0.44% Triton X-100) in a 1:9 (v/v) proportion under strong magnetic agitation, resulting in the substrate solution. After, the supernatants (10 μL) were mixed with the substrate solution (90 μL) and read immediately at 410 nm (time zero). The mixture was incubated at 37 °C and read again after 30 min in SpectraMax spectrophotometer (Molecular Devices, USA). As a control, buffer was added instead secreted extract. Results were obtained in μmol of ρNPP per hour.

Protease activity was evaluated using 2 mM of subtilisin (Pr1) substrate (N-suc-ala-ala-pro-phe-ρNA) and trypsin (Pr2) substrate (Bz phe-val-arg-ρNA) in 100% DMSO. Approximately 2 μg of protein of each supernatant was added to 50 mM Tris-HCl pH 8.0 buffer to complete 100 μL. Kinetic assays were monitored at 37 °C for 30 min in a SpectraMax. One protease unit (U) was defined as the amount of enzyme that produces one ρmol of ρ-nitroaniline per hour, in the assay conditions described.

For catalase activity, hydrogen peroxide was used as substrate and phosphate buffer was added along with the substrate at 10 mM to 25 μL sample aliquots. The reduction in hydrogen peroxide concentration was tracked by measuring the decrease in absorbance at 240 nm over a period of three minutes (E = 39.4 mM cm^-1^).

For phospholipase C (PLC) activity, ρ-nitrophenylphosphorylcholine (ρNPPC) was used, which is a chromogenic substrate selectively hydrolyzed by PLC. The ρNPPC was prepared at a 20 mM concentration in a 50 mM Tris-HCl pH 8.0 and 60% sorbitol buffer. Samples (10 μL) were mixed with 90 μL of substrate solution and incubated at 37 °C for 1 hour. Control was made with buffer instead sample. After this time, the assay was read at 410 nm in SpectraMax spectrophotometer. One unit of PLC was defined as described above for lipolytic activity unit.

For phosphatase activity, the rate of ρ-nitrophenol (ρ-NP) production was assessed according to (101). Samples were incubated for 1 hour at room temperature in 0.2 mL of reaction mixture (116.0 mM NaCl, 5.4 mM KCl, 30.0 mM Hepes-Tris buffer pH 7.0, and 5.0 mM ρ-NPP). Enzymatic activity was triggered by adding the samples and halted with 0.2 mL of 20% trichloroacetic acid. Then, an aliquot of 0.1 mL of the supernatant was added to a 96-well plate, along with 0.1 mL of NaOH. Readings were made at 405 nm using the SpectraMax spectrophotometer. The enzymatic activity was then calculated by subtracting the non-specific ρ-NPP hydrolysis, measured for control (made with buffer), and the concentration of ρ-NP was estimated using a standard calibration curve.

### Bioassays

For the bioassays, engorged *R. microplus* females from infested bovines were disinfected using hypochlorite 2.5% for 2 s, washed with sterile saline and sterile distilled water (102). Then, ticks were completely immersed in a 10^8^ conidia mL^-1^ suspension of E6 or NE strains for 15 s. After exposure, ticks were sorted into three independent groups of 10 individuals in Petri dishes and kept in a humid chamber (> 90% relative humidity) at 28 °C. They were inspected to check for the surviving animals every day. Controls were immersed in sterile distilled water instead spore suspensions, receiving the same conditions described here.

### Proteome analysis

#### Sample preparation for mass spectrometry

For secretomic analysis, supernatants of each strain were promptly boiled for 5 min, in order to inactivate endogenous proteases that could affect the secretome results. Then, the samples were lyophilized and stored at −80 °C until use. Then, 100 μg of proteins were resuspended in digestion buffer (8 M urea, 100 mM Tris-HCl, pH 8.5) and digested with trypsin (2 μg) (Promega, Madison, WI) for 16 h at 37 °C as previously described (75). The reaction was then stopped by adding 5% formic acid.

After digestion, the proteins were pressure-loaded into a biphasic capillary column containing 2.5 cm ion exchange resin (Partisphere SCX) and 2 cm reverse phase resin (Acqua C18). MudPIT salt separation included twelve steps, utilizing a gradient ranging from 0 to 100% of buffer B (80% acetonitrile/0.1% formic acid) (103). Peptide samples were then loaded onto a LTQ-XL system (Thermo Fisher, USA), following the manufacturer’s protocol. Each analysis cycle consisted of one full-scan mass spectrum (300-2000 m/z) followed by five data-dependent MS/MS spectra at a 35% normalized collision energy was repeated continuously throughout each step of the multidimensional separation. Dynamic exclusion was enabled with a repeat count of 1, a repeat duration of 30 s, and an exclusion list size of 200, as previously described (75) in order to prevent repetitive analysis. The Xcalibur data system (Thermo, CA) was used to control both the mass spectrometer scanning and the HPLC solvent gradient functions. Mass spectrometry analysis was performed in six different runs per condition with each one prepared from an independent biological replicate/liquid culture.

### Protein identification, quantification, and molecular characterization of the differential secretome

Different software programs were used to characterize the secretomes obtained from mycelia and conidia of both strains, leading to an evaluation of the molecular and functional properties of the secretomes. PatternLab V (104) was used in the identification of the proteins, providing a relation of which were exclusively identified in each condition (AAPV module) and which were differentially expressed (TFold module), along with the up and down regulated proteins for each growth condition. Protein identification was based on the *M. anisopliae* strains E6 and NE sequenced genomes described in this article. Redundant protein sequences of the merged genomes used as database were removed with the SeqKit2 package within the R environment, using the seqkit function (105). The parameters used were: proteins detected in at least four out of six replicates per condition, t-test with a p-value of 0.005, and BH q-value of 0.05 (5% FDR). The BlastP suite (https://blast.ncbi.nlm.nih.gov/Blast.cgi) was used in order to further verify every detected protein sequence, check hypothetical proteins, and re-annotate those with enough corresponding matches or identify conserved domains.

The STRING Platform (https://string-db.org), providing information of functional protein association networks was used to categorize the differentially expressed proteins (DEPs) (p-values < 0.05) and Gene Ontology (GO) terms related to biological processes (BPs) and molecular functions (MFs).

The search for conserved secretion signal prediction was performed using four online programs: Phobius (https://phobius.sbc.su.se/), PrediSi (http://www.predisi.de/), SignalP 6.0 (https://services.healthtech.dtu.dk/services/SignalP-6.0/) and TargetP 2.0 (https://services.healthtech.dtu.dk/services/TargetP-2.0/). A protein was considered to have a positive prediction for a secretion signal if at least two out of the four programs yielded a positive result.

### Bioinformatic analysis

The detailed scripts of the bioinformatic analysis are available here: <u>michaelH-git/StarshipTE_Metarhizium_anisopliae</u>

In short:

Assembly. Assemblies were generated from PacBio Hifi Reads using Canu (version 2.2) (106) and the assemblies were visually inspected using Tapestry (107) and contigs that showed less than 20% or more than 300% of the average genome-wide coverage, as calculated by mosdepth (version 0.3.4) (108) on the minimap2 (version 2.26-r1175) (109) aligned PacBio reads were removed, because these were considered to be mitochondrial genome, spurious contigs or contamination. The resulting assemblies were scaffolded using the PacBio reads by ntLink (version: v1.3.11) (110) which led to seven contigs and three scaffolds, and two contigs to one scaffold in strain E6, leaving all other contigs unmodified. These final assemblies were again visually inspected by Tapestry and telomeric sequences (TTAGGG) were detected. The general assembly statistics are available in TableS1. For E6 all scaffolds have telomeric repeats at both ends, therefore representing chromosomes, while in NE three scaffolds have telomers at both ends, three have telomers only on one end, and two have no telomeric sequences. The mitochondrial genomes were identified by MitoHifi (version: v3.2.2)(111).

Small (accessory) chromosomes were identified by extracting DNA from chromosomal bands separated by PFGE as described in (43). The extracted DNA was then sequenced using a low-input DNA library preparation protocol at the Max Planck Sequencing Center in Cologne on an Illumina NextSeq2000 platform. Reads were trimmed using Trimmomatic (112) and quality was inspected using FastQC (version v0.12.1) (113) and mapped to the respective genome assemblies using Bowtie2 (version 2.5.1) (114). Duplicates were marked, and read groups were added using Picard. Sequencing coverage in 50 kb windows was calculated using mosdepth (version 0.3.4) (108).

The genome assemblies were annotated for transposable elements (TEs), genes, secondary metabolite cluster candidates, functional gene annotation, BUSCO completeness (against the *ascomycota_odb10* dataset), tRNA repertoire, gene-wise relative synonymous codon usage (RSCU), potentially secreted proteins, effector candidates, and CAZymes. The annotation was performed using a combination of tools: EDTA (version 2.2.2) (78), BRAKER3 (version 3.0.8) (115) with fungal protein information from the OrthoDB11 database as evidence, antiSMASH (version 7.1.0) (116), BUSCO (version 5.7.1) (117), eggNOG-mapper (version 2.1.12) (118), and InterProScan (119) using the interproscan-5.68-100.0 database. tRNA genes were identified using tRNAscan-SE (version 2.0.12) (120), and functional gene annotation was further supported by BLAST (121) searches against the SwissProt database restricted to fungi. Codon usage and RSCU values were calculated using bioKIT (version 1.1.3) (122), and CAZymes were annotated with dbCAN (123). Functional annotations were collated and visualized using Blast2GO (version 6.0.3) (124). Starships were identified among 19 *Metarhizium* sp. genomes and annotated using Starfish (79). Starships were visually confirmed and only Starships with at least 50000 bp total flank-alignment were kept. Phylogenetic trees were constructed on MAFFT alignments using IQ-TREE2 (125) with 10000 bootstrapping and automatic model detection. Phylogenetic trees were visualized using iTOL (126).

To determine the SNPs between NE and E6 the E6 PacBio reads were mapped onto the NE genome assembly minimap2 (version 2.26-r1175) (109), and subsequent SNP calling and file processing were performed with SAMtools (version 1.17) (127), BCFtools (version 1.17) (127), and BEDTools (version 2.31.0) (128).

Orthogroups were defined using OrthoFinder (version 2.5.5) (129), and synteny based on orthologous genes was determined with GENESPACE (version 1.3.1) (130). Syntenic blocks were visually inspected, and adjacent syntenic regions with identical orientation were manually merged if they were not separated by more than 5 gene models. Breakpoint regions were defined as the boundaries between adjacent regions located on different syntenic chromosomes and/or with differing orientations. For each identified breakpoint, the region in the *M. anisopliae* NE genome plus 25 kb of upstream and downstream flanking sequence was extracted and aligned to the corresponding syntenic region in *M. anisopliae* E6 using MUMmer (131), as implemented in pyGenomeViz (version 1.5.0) (https://github.com/moshi4/pyGenomeViz/), and visualized with the same tool. Breakpoint coordinates were extracted from the MUMmer alignment output. Only those breakpoint regions where visual inspection allowed the localization of the breakpoint within a 10 bp window were retained. These 10 bp sequences which contained the breakpoints were subsequently aligned with each other using MAFFT (version 1.5.0), as implemented in Geneious Prime (version 2023.1.2).

A total of 632 Starships were downloaded from Starbase (77) on 10.03.2025. After comparison and visual inspection, 618 Starships were determined to be likely unique. Please note: The accessions on Starbase might change upon its final release. Hence, we provide all information using the preliminary accessions in Table S4 and in addition the fasta file containing the sequences in Supplementary File S1. For those Starships for which the genome accession was not available, the genome assembly information was retrieved from NCBI for 383 Starships using the genomic location IDs provided by Starbase. Transposable elements (TEs) were annotated on all Starships (retrieved from Starbase and those identified in this study) using EDTA, including a Helitron consensus sequence identified as having recently increased in copy number in *M. anisopliae* strain NE. This Helitron sequence was included because of the low performance of EDTA to annotate it. Only repetitive regions longer than 100 bp and assigned to known TE families were considered for further analysis; repetitive regions lacking a TE classification were excluded. FASTA sequences of annotated TEs were extracted and subjected to BLAST searches against (i) their genome of origin—where all Starship regions had been hard-masked—and (ii) all Starships. BLAST hits were filtered to retain only those with >99% sequence identity and >99% coverage of the query. These were further categorized to identify hits with 100% identity and coverage. For each Starship, the number of TEs with at least one qualifying BLAST hit was recorded. Since similar or identical TEs and subsequences often exist within the same Starship, raw counts of BLAST target hits would be inflated, hence Blast target hits were merged if they overlapped >99% (of the smaller sequence) before counting. To test for the overrepresentation of perfect TE copies (100% sequence identity covering 100% of the length) on Starships, the number of perfect copies found within Starships was compared to those found outside, adjusting for the respective sizes of Starship and non-Starship regions. Statistical significance was assessed using Fisher’s exact test and corrected for multiple comparisons using the conservative Bonferroni adjustment. The genome accessions for publicly available genome assemblies of the *Metarhizium* species are described in Table S7, assemblies for published Starships are described in Table S4.

Note: While scripting for the parallelization of bioinformatics procedures was aided by LLMs (ChatGPT 4.0), the entire bioinformatics procedure, including the choice and setting of tools and commands, remained the sole responsibility of the authors.

## Data availability

The sequencing reads, genome assemblies of *M. anisopliae* strain NE and strain E6 were deposited on NCBI using the bioproject PRJNA1277033 and PRJNA1277034.

## Acknowledgements

We particularly would like to thank Adrian Forsythe, Aaron Vogan and Emile Gluck-Thaler for providing the extremely helpful Starbase database, without much of the analysis would not be possible.

## Supplementary Tables

Table S1: Overview of Assembled Genomes for *M. anisopliae* strain NE and E6 used in this study. Table S2A: Syntenic regions based on orthologous genes.

Table S2B: Locations of major breakpoint regions in *M. anisopliae* strain NE and their association with the three expanded TE families.

Table S3A: Overview of Starships identified in *Metarhizium* species in this study. Table S3B: Overview of tyrosine recombinases in *Metarhizium* Starships.

Table S4: Characteristics and TE content of published unique Starships and Starships identified in *Metarhizium* species in this study.

Table S5: Results of differential Secretome Proteomics comparing *M. anisopliae* strain E6 with NE and co-localization of genes with structural variation in *M. anisopliae* strain NE.

Table S6: Results of BLAST of TE sequences located on a Starships against all Starship sequences. Only BLAST hits of at least 99% sequence identity and 99% sequence coverage are included.

Table S7: Accessions of published *Metarhizium* and *Epichloë festucae* genome assemblies used in this study.

## Supplementary Files

Supplementary File S1: FASTA sequences of all Starships used in this study. These include those identified in Metarhizium in this study (39) and those downloaded from Starbase (618).

## Supplementary Figures

**Fig. S1:**
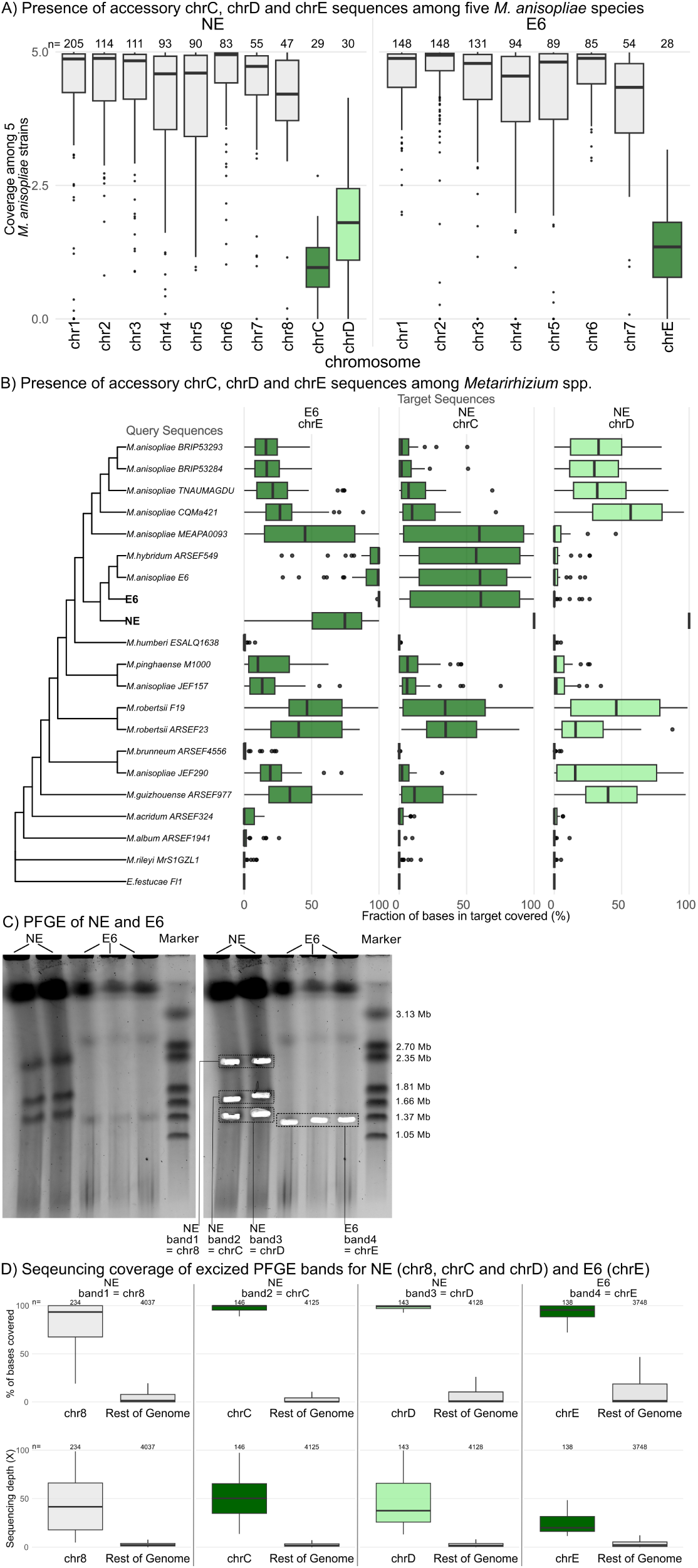
*M. anisopliae* strain NE contains two accessory chromosomes (chrC and chrD) and one additional core chromosome (chr8), while strain E6 contains one accessory chromosome (chrE). **A)** Accessory chromosomes were identified based on presence-absence polymorphisms across five previously published *M. anisopliae* isolates. The data show sequence coverage, defined as the number of assemblies (out of five) in which a given 50 kb window (excluding transposable elements) is present. A coverage value of 5 indicates presence in all five isolates. Core chromosomal sequences (grey) display a median coverage near 5, while two entire chromosomes in strain NE (green and light green) and one in strain E6 (green) show substantially lower coverage, suggesting they are not shared among all isolates and can therefore be considered accessory. **B)** The presence of accessory chromosomes chrC, chrD, and chrE was evaluated across 19 published genomes of *Metarhizium* species and *Epichloë festuciae* (used as an outgroup), as well as in the NE and E6 strains characterized in this study. The proportion of the accessory chromosome sequences (in 50 kb windows, excluding transposable elements) covered by each genome is shown as boxplots on the right, with a cladogram on the left depicting the phylogenetic relationships among genomes (based on 4560 single-copy orthologues; see Fig. 2). **C)** Accessory chromosomes and core chromosome 8 were also identified by PFGE (pulsed-field gel electrophoresis) of strains NE and E6, shown before (left) and after (right) excision of chromosome bands. Excised bands were labelled, pooled as indicated, sequenced using Illumina reads and mapped to long-read assemblies to identify specific chromosomes. **D)** Boxplots show the percentage of base coverage (top row) and sequencing depth (bottom row) of Illumina reads for identified chromosomes (green), compared to the rest of the genome (grey), calculated in 10 kb windows.

**Fig. S2:**
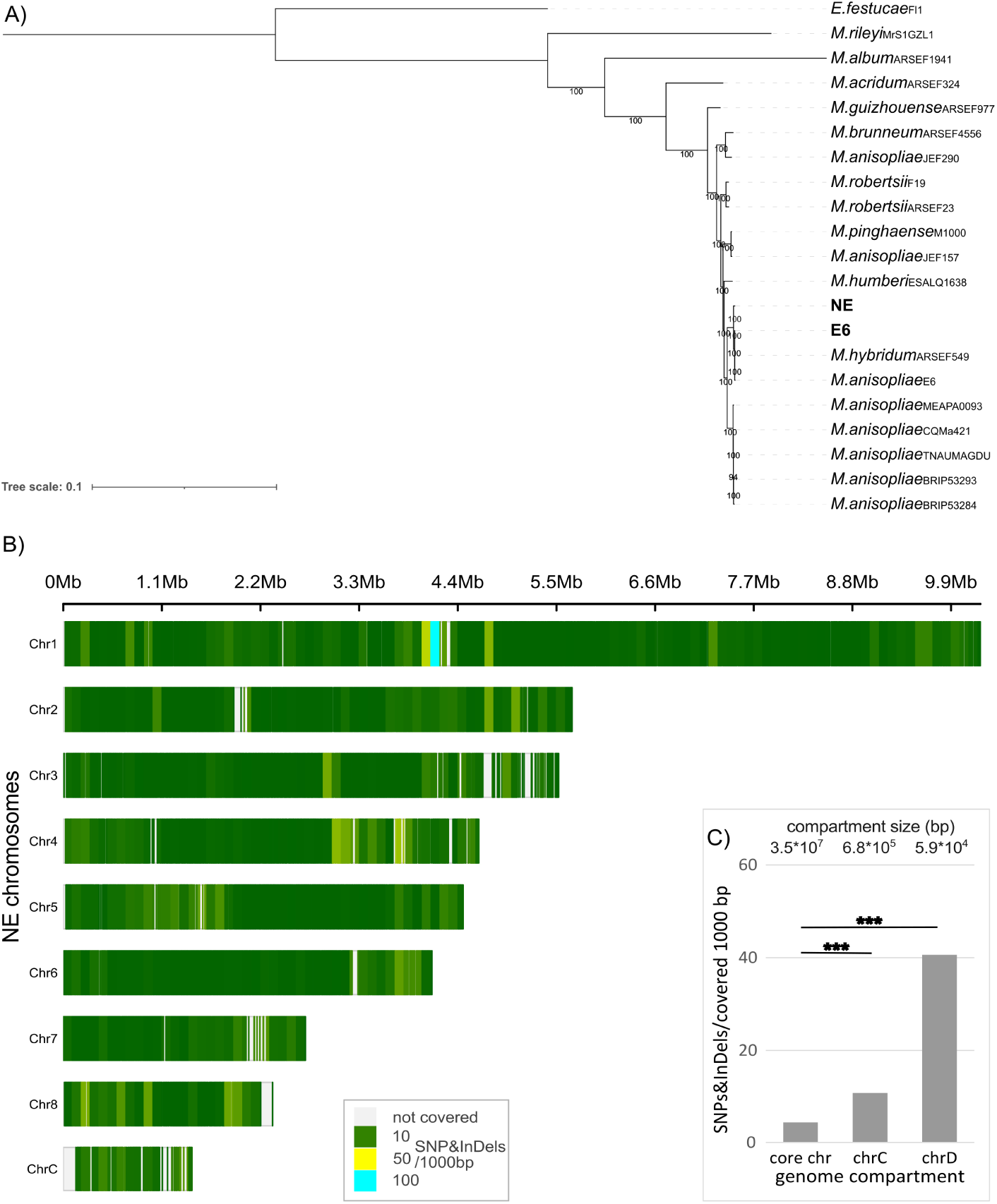
Phylogenetic and SNP/INDEL analyses confirm a close evolutionary relationship between strains NE and E6, with no evidence that the mosaic syntenic regions on NE chromosomes evolved along distinct trajectories. **A)** Phylogenetic analysis was performed using concatenated protein sequences from 4,383 single-copy orthologues across 20 *Metarhizium* isolates, with *Epichloë festuciae* used as an outgroup. Bootstrap support values from 1,000 replicates are shown as percentages. *M. anisopliae* strains NE and E6 are closely related and form a monophyletic clade that includes a previously published E6 assembly. **B, C)** Comparison of SNP and INDEL densities across core and accessory chromosomes, calculated per 1,000 base pairs of covered sequence (excluding transposable elements). Core chromosomes exhibit low variation, except at the rRNA gene cluster on chromosome 1. This variation does not correspond to the mosaic of different syntenic E6 chromosomes along the NE chromosomes. In contrast, accessory chromosomes chrC and chrD exhibit higher SNP and INDEL densities, although only 58,529 bp are covered in these regions. Statistical significance, determined using Fisher’s exact test, is indicated by *** (p < 0.001). Note: In panel B, scaffolds corresponding to chrD and scaffold_15 are omitted for clarity because there were only limited syntenic regions between E6 and NE on these sequences.

**Fig. S3:**
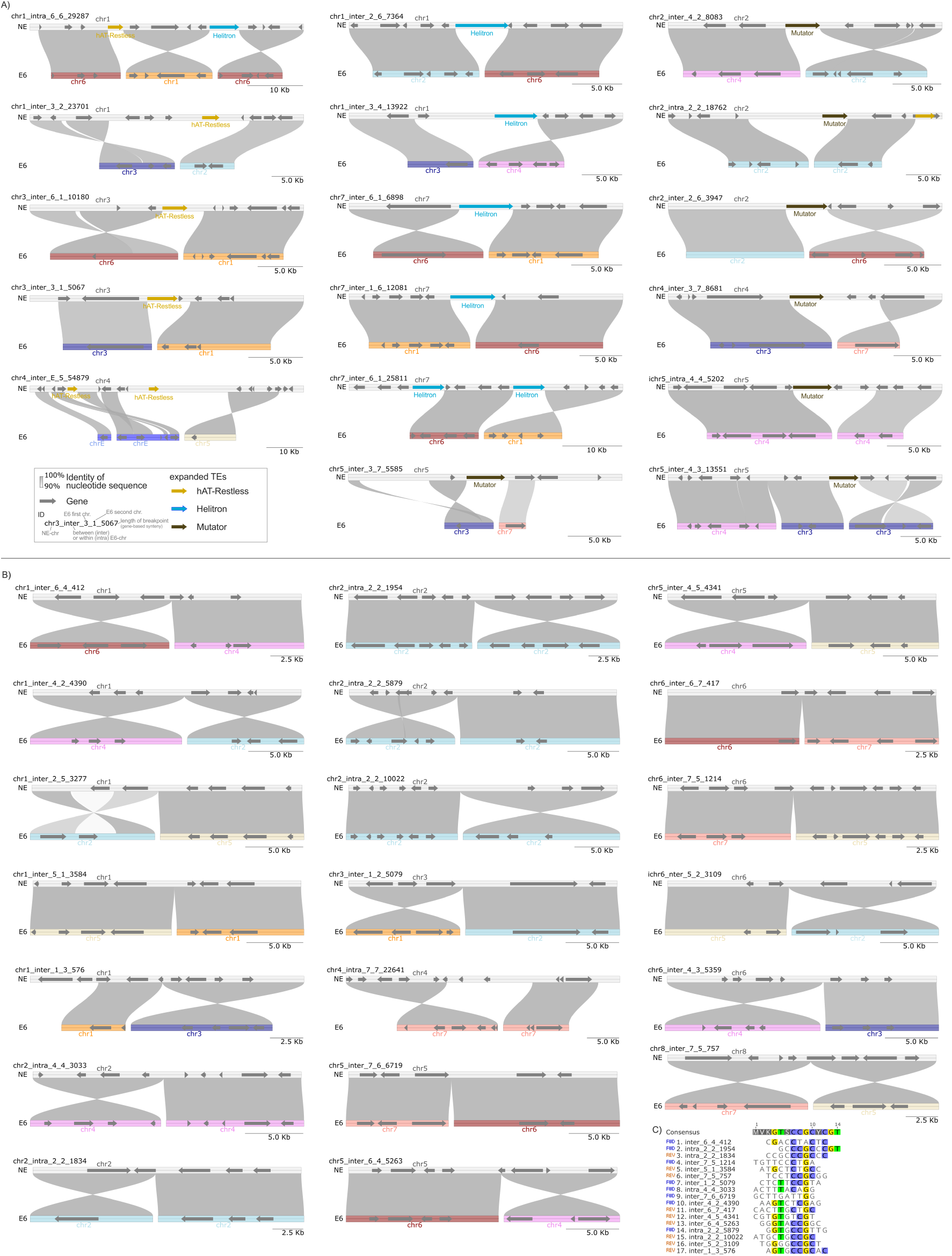
Nucleotide synteny at breakpoints regions where (A) expanded TEs are present or (B) not present. Breakpoint regions were defined by macrosynteny analysis based on gene synteny, and nucleotide-level synteny at these breakpoints was analyzed using MUMmer. **A)** A total of 17 breakpoints are associated with the three expanded TE families, often suggesting recombination occurred at or within the expanded TE families. **B)** A total of 20 breakpoint regions are not associated with expanded TEs; in these cases, breakpoints appear to result from direct recombination between syntenic sequences. For 17 of these breakpoint regions, the precise breakpoint could be identified within a 10 bp region. **C)** MAFFT alignment of the 10 bp regions flanking the 17 identified breakpoints, with bases that match the consensus sequence highlighted for comparison.

**Fig S4:**
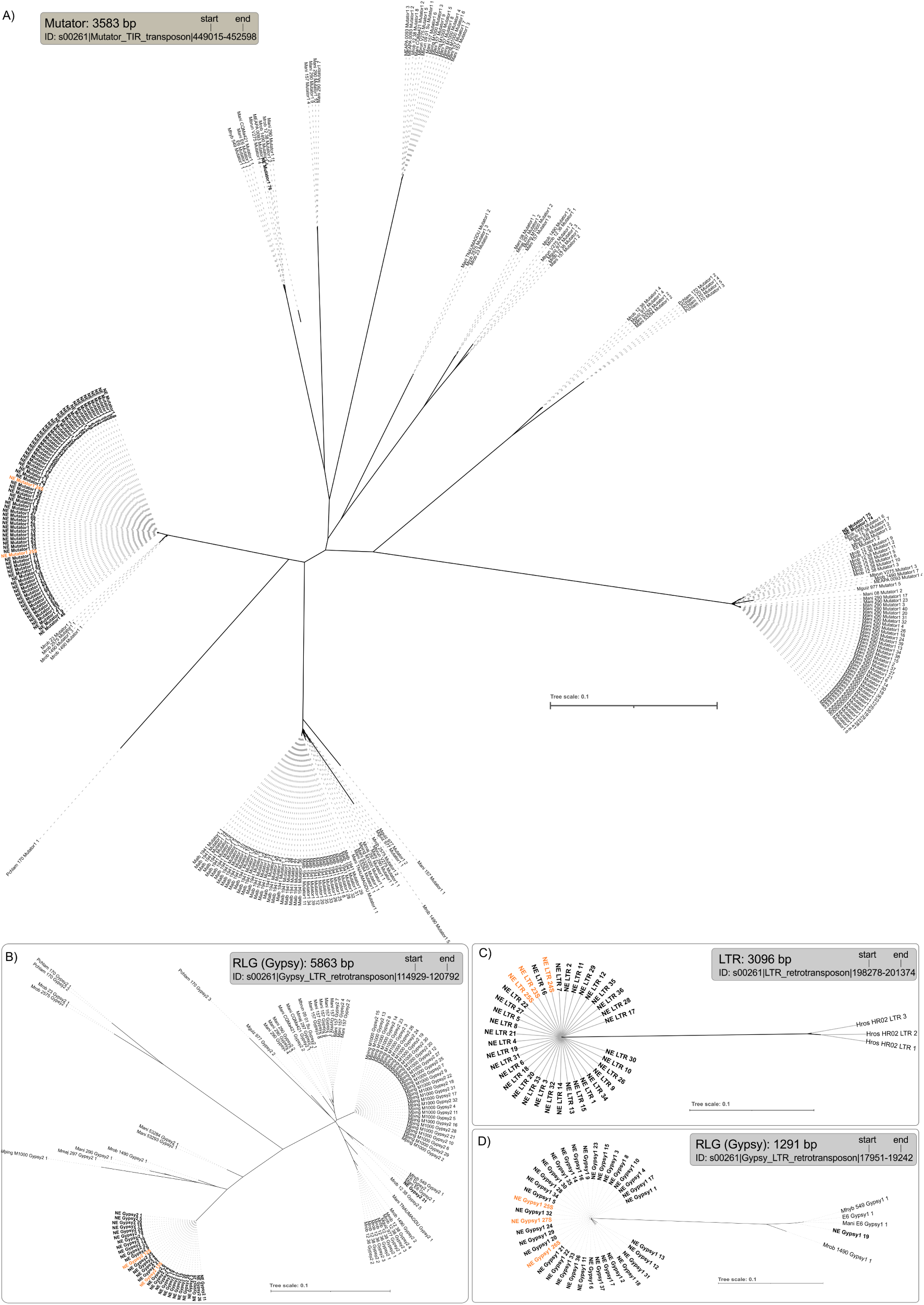
Phylogenies of transposable elements (TEs) on Starship S000261 and expanded in *M. anisopliae* strain NE. TEs expanded in NE are shown in bold, with those located on Starship S000261 highlighted in orange. The figure includes phylogenies of the following TEs: **A)** an expanded Mutator TE associated with structural variation, **B)** an expanded RLG (Gypsy) TE, **C)** an expanded LTR TE, **D)** a second expanded distinct RLG (Gypsy) TE. The ID and the location within the Starship of the sequences used to identify additional copies are given above the phylogenies. The analyses reveal that the TEs on the Starship form a monophyletic group with the expanded TEs found elsewhere in the genome and are distinct from other copies of the same TE type, suggesting a separate origin. Consensus trees were generated from 1,000 bootstrap replicates of MAFFT-aligned sequences using IQ-TREE with TIM+F+G4 as the best-fit model. Species abbreviations are provided in Table S6, and, in addition to the *Metarhizium* species, *Pochonia chlamydosporia* strain 170 (Pchlam 170) and *Hirsutella rhossiliensis* HR02 (Hros HR02) were included as outgroups. Sequences showing 80% sequence identity and 80% coverage to the query sequence are displayed. Note: For the LTR, since no hits were obtained within the *Metarhizium* genus, we included sequences with at least 60% identity and coverage.

**Fig. S5:**
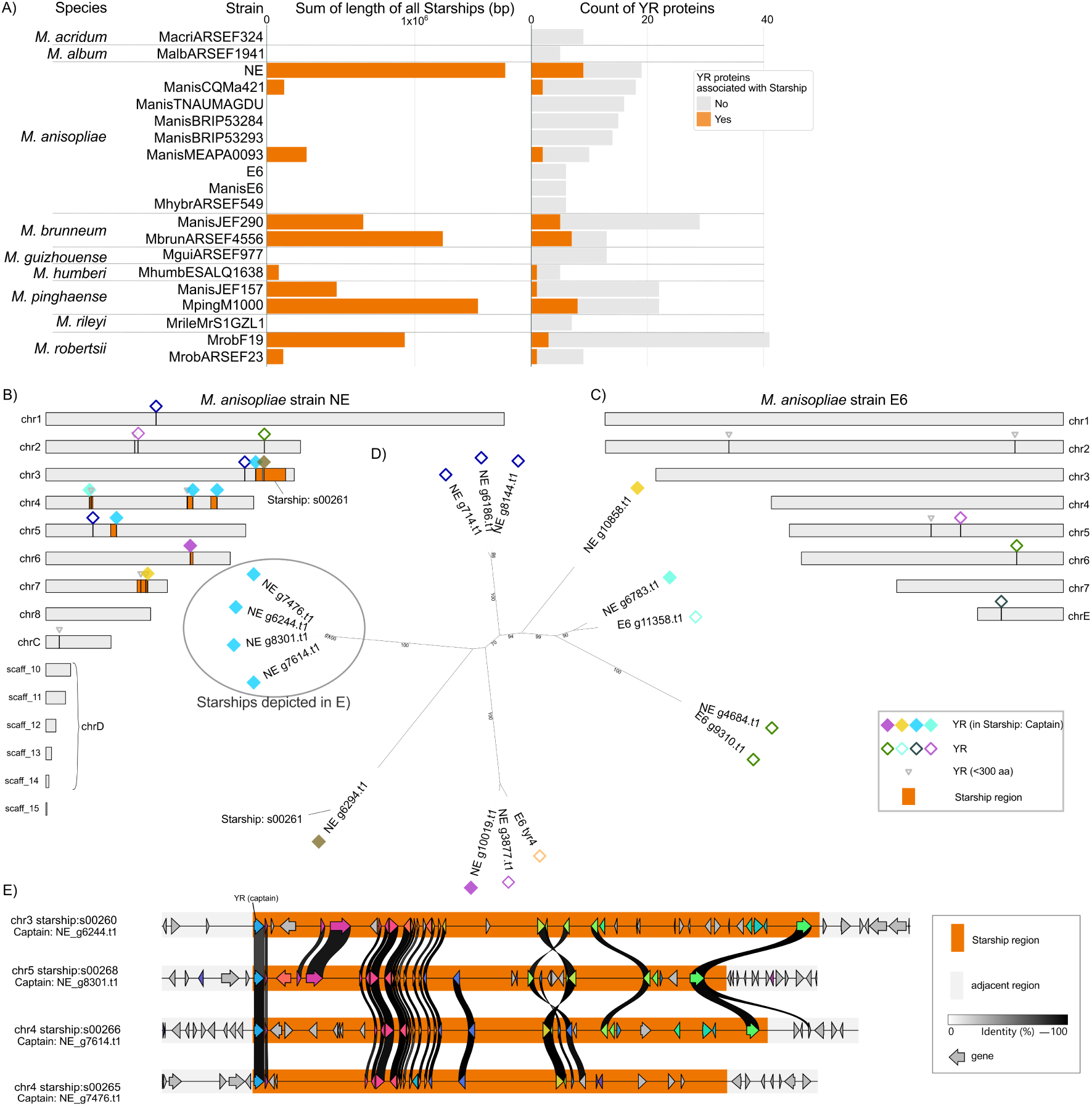
Distribution patterns and evolutionary relationships of Starships across *Metarhizium* species. A) Total length of Starships identified in each indicated species (left), and the number of tyrosine recombinase (YR)–encoding genes, either associated with active Starships (i.e., Captains, shown in orange) or not (shown in grey), is shown on the right. High numbers of YR-encoding genes were detected in several *Metarhizium* species, with particularly high numbers of Captains in *M. anisopliae* strain NE, *M. brunneum*, and *M. pinghaense*. B–C) Comparative karyograms of *M. anisopliae* strains NE and E6, with Starship insertions differentiating the two highlighted in orange. Multiple Starships were observed in the NE strain, which harbors eight such insertions, whereas the E6 strain lacks such elements. D) A consensus tree based on YR proteins (>300 amino acids) reveals a close relationship among four YR proteins associated with Starships in the NE strain (highlighted in pink). E) Synteny analysis of the four Starship elements and their flanking regions shows a subset of highly conserved cargo gene content but distinct insertion sites, with no detectable synteny. This pattern suggests that these elements represent independent invasion events rather than duplications of a single ancestral element. Note for panel E: Genes are displayed, with paralogous genes shown in the same color.

**Fig. S6:**
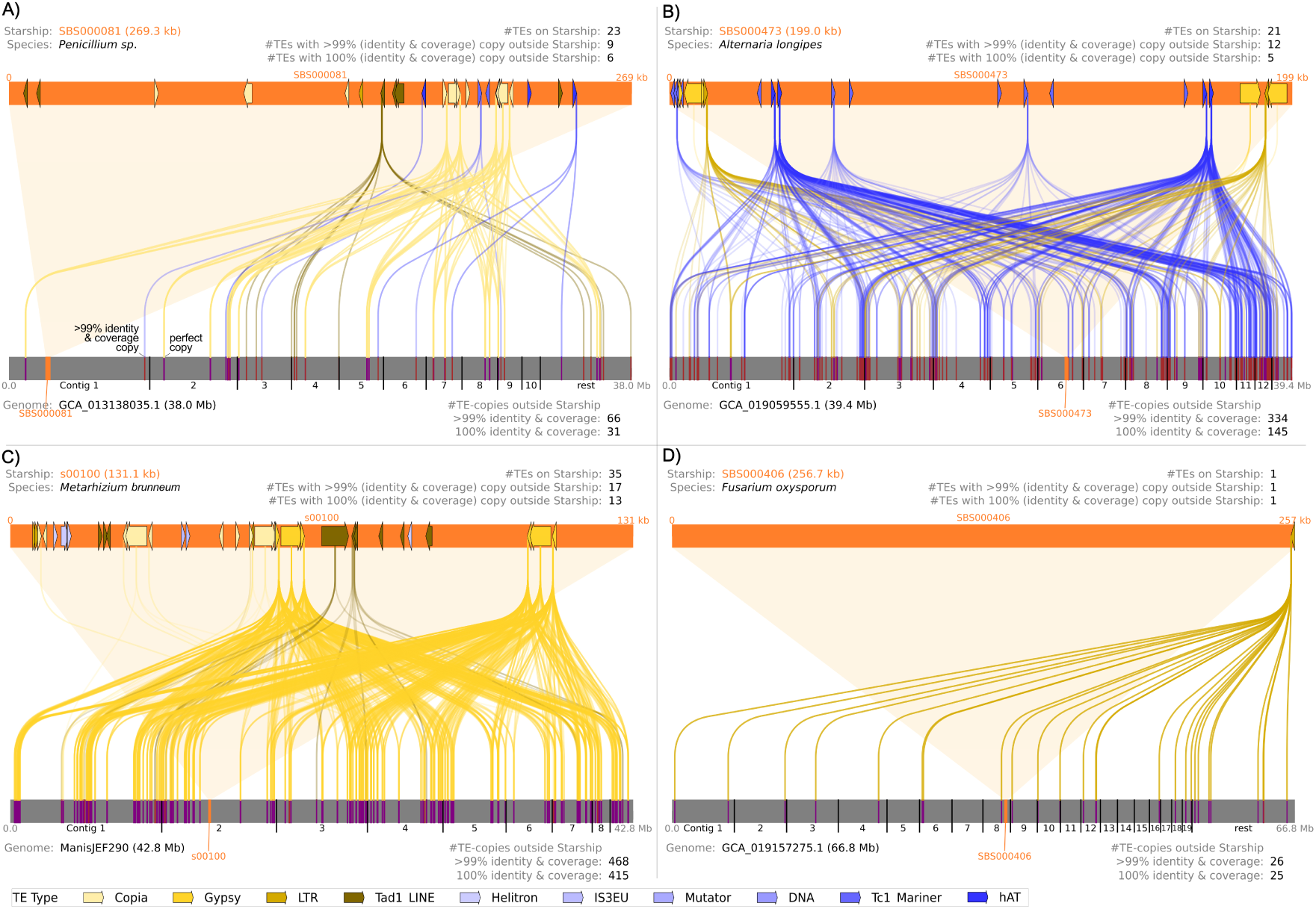
Examples of various TE expansion from different Starships. The type and extent of transposable element (TE) expansions vary between different Starships. TE distributions are shown for four representative Starships (orange bars at top) and their perfect (100% identity and coverage; purple markers) or highly similar (>99% identity and coverage; brown markers) genomic copies outside the Starships (continuous grey genome bars below). These examples illustrate that many TEs found on Starships have numerous identical or highly similar copies elsewhere in the genome and that different TE families can be expanded from the same Starship. Examples shown are A) SBS000081 in *Penicillium sp.,* B) SBS000473 in *Alternaria longipes*, C) SBS00100 in *Metarhizium brunneum*, and D) SBS000406 in *Fusarium oxysporum*. TE families are color-coded: Retrotransposons appear in different shades of yellow, while DNA transposons are shown in various shades of blue. Note for panel C: the isolate ManisJEF290 was classified as *M. brunneum* based on the phylogenetic analysis presented in this study.

**Fig. S7:**
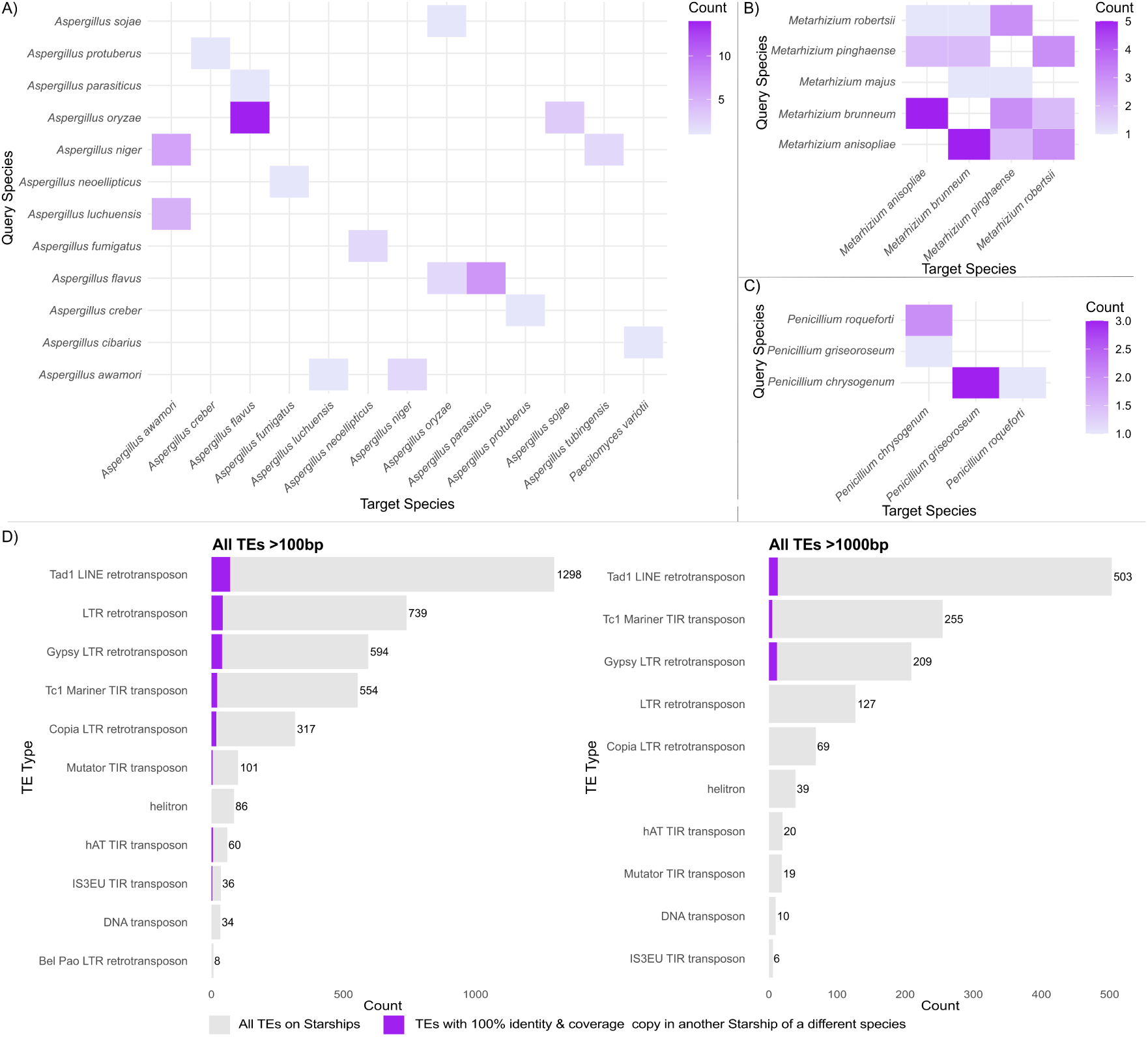
Transposable elements (TEs) on Starships appear to be predominantly horizontally transferred within genera, although there are also some examples of transfers occurring between different genera. A–C) Heatmaps illustrating the number of Starships containing at least one TE that has an identical copy (100% sequence identity and 100% coverage) in at least one Starship from a different species, shown separately for *Aspergillus*, *Metarhizium*, and *Penicillium*, respectively. TEs from query Starships were blasted against Starships of target species, and the number of Starships containing perfect TE copies in the target species was counted. Within the genus *Aspergillus*, many identical TE copies are shared between *A. flavus* and *A. oryzae*; in the genus *Metarhizium*, TEs are commonly shared between Starships from *M. anisopliae* and *M. brunneum*; and in the genus *Penicillium*, perfect TE copies frequently occur between *P. chrysogenum* and *P. griseoroseum*. Although perfect TE copies are almost exclusively shared between Starships within the same genus, one notable exception involves a 561 bp Mutator transposon located on a Starship from *A. cibarius*, which shares an identical copy with a Starship from *Paecilomyces variotii*, belonging to a different family. Another example of intergeneric transfer involves a 400 bpTC1-Mariner TE shared between Starships found in *Exserohilum turcicum* and *Bipolaris maydis* (see Table S6). D) A total of 3,827 TEs (of which 1,257 are longer than 1,000 bp), belonging to various TE families, were identified on Starships. Of these, 211 TEs (including 30 longer than 1,000 bp) have at least one perfect copy on a Starship from a different species. No clear preference is observed for specific TE families to be shared between Starships of different species.

**Fig. S8:**
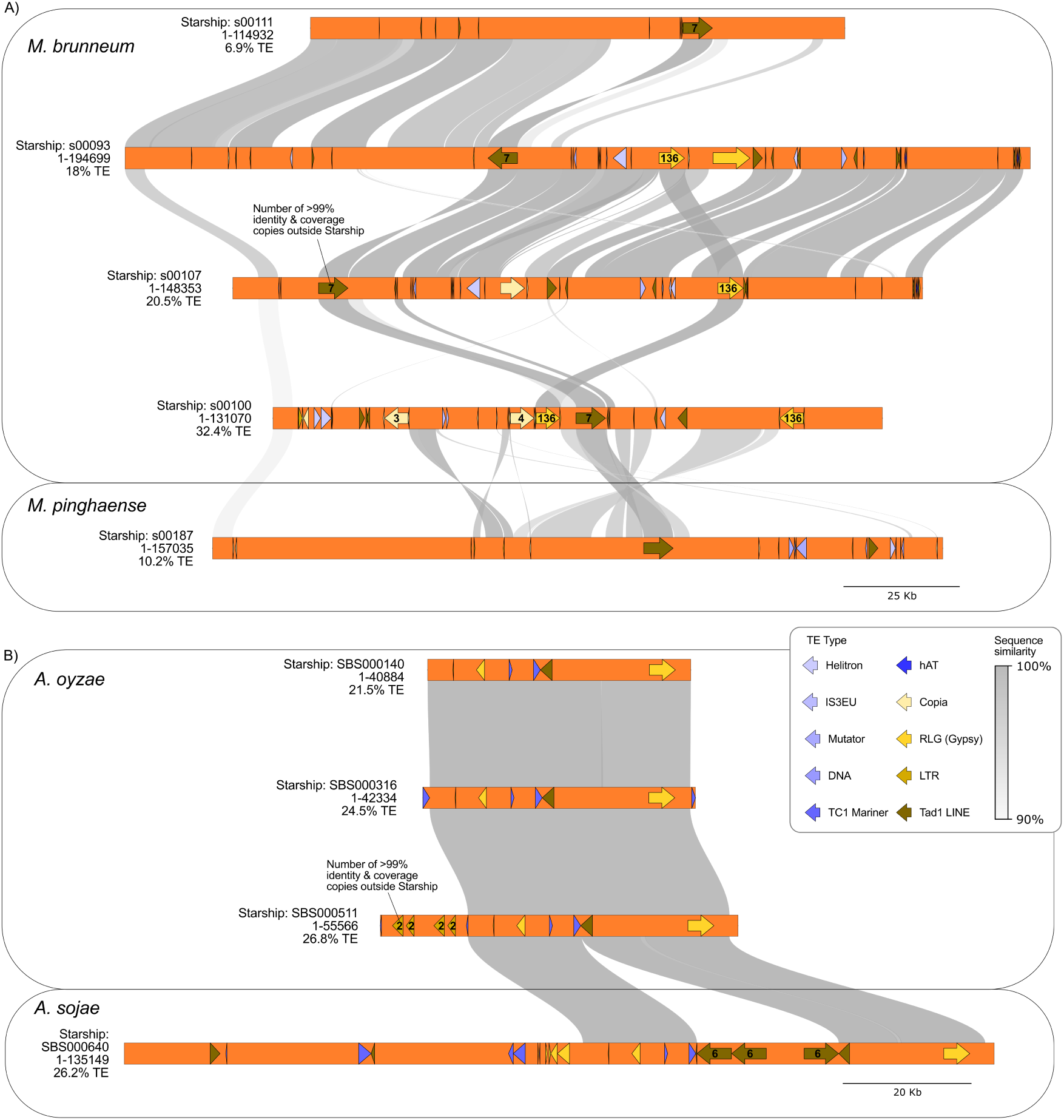
Starships can act as both the source and destination of transposing transposable elements (TEs), facilitating their movement between species. A) An example of TEs shared among five different Starships (in orange), with the upper four from *M. brunneum* strain JEF290 and the lowest from *M. pinghaense*. For clarity, only TEs are indicated by arrows (genes are omitted). A Tad1 LINE transposon, among other sequences, is shared between Starship s00187 from *M. pinghaense* and Starship s00100 from *M. brunneum*. This element, along with others, has expanded in copy number across the genome of *M. brunneum* and most likely also invaded three additional Starships (s00111, s00093, and s00100). Synteny breaks and altered TE orientations relative to conserved Starship structure provide evidence for recent TE movement. B) An example of TEs shared across species and the acquisition of new TEs onto Starships is shown with four Starships from two species—three (on top) from *A. oryzae* and one from *A. sojae*. Large syntenic regions across all four Starships containing TEs indicate interspecies TE transfer via Starships. In SBS00511 of *A. oryzae* and SBS00640 of *A. sojae*, TEs that have expanded throughout the genome appear to have recently invaded the Starships. Numbers in the TEs indicate the count of copies with >99% sequence identity and coverage found outside Starships. Synteny breaks involving these TEs, compared to the surrounding syntenic Starship regions, further support recent transposition events. For clarity, only TEs longer than 1,000 bp are labeled with copy numbers. TE families are color-coded: Retrotransposons appear in different shades of yellow, while DNA transposons are shown in various shades of blue. Sequence identity is visualized by connectors in different shades of grey.

## Notes

### Competing Interest Statement

The authors have declared no competing interest.

